# On the Genetic Origins of Phenotypes in Genome-Wide Association Studies: The SNP Allocation For Estimating Heritability (SAFE-*h*^2^) Tool to Explore Additive-only Allelic Effects or Additive and Non-Additive Allelic Effects

**DOI:** 10.1101/2023.08.28.555092

**Authors:** Behrooz Darbani, Mogens Nicolaisen

## Abstract

Polygenicity requires the inclusion of all SNPs to ensure accurate estimation of SNP heritability. However, SNPs with large association *p*-values should be excluded, as they impose biologically meaningless negative contributions, *i.e.*, downward bias, to heritability estimates. This is demonstrated with simulated data and validated using real-world datasets. We introduce SAFE-*h*^2^ as a solution to this dilemma. SAFE-*h*^2^ performs heritability profiling and establishes a *p*-value threshold to filter out SNPs with negative contributions. By examining 74 phenotypes across eight species, SAFE-*h*^2^ revealed an average negative contribution of 30 units, imposed by SNPs from the upper-bounds of *p*-value thresholds. SAFE-*h*^2^ also provides a safe *p*-value interval to minimize false-positive SNP hits and facilitates capturing intra-locus (*Ref* -allele_SNPi_ × *Ref* -allele_SNPi_, *Ref* - allele_SNPi_ × *Alt* -allele_SNPi_, and *Alt* -allele_SNPi_ × *Alt* -allele_SNPi_) additive and non-additive effects within the framework of linear models. The allelic adjustment algorithm of SAFE-*h*^2^ revealed considerable uncaptured phenotypic variance when relying on additive effects and improved the SNP heritability estimations by up to 56 units across 50 phenotypes in six species through the combined capture of both additive and non-additive allelic effects.

## Introduction

Single nucleotide polymorphism (SNP) heritability is a key measure in genome-wide association studies (GWAS), reflecting the experimental power for exploitation of phenotypic variance. Estimating heritability requires both genetic variance (**σ_g_^2^**) and phenotypic, *i.e*., total, variance (**σ_p_^2^**), with **σ_g_^2^** ≤ **σ_p_^2^** and heritability (**σ_g_^2^** ÷ **σ_p_^2^**) values ranging from 0 to 1 (0–100%). Compared to broad sense heritability (*H* ^2^) estimates, SNP heritability is typically estimated on additive effects and is thus considered as narrow-sense heritability^1,2^. Nevertheless, SNP heritability provides valuable insights into the genetic architecture of traits and has significant applications in plant and animal breeding, as well as medicine. SNP heritability is a critical metric not only for guiding future research and calculating response to genomic-selection but also for assessing the genetic influence on the trait-of-interest in genome-wide association studies, as reporting significant SNP associations alone is insufficient. Therefore, accurate estimation of SNP heritability is crucial, as both underestimation and overestimation can lead to substantial genetic misinterpretations. For instance, a significant underestimation of SNP heritability for a trait, *i.e*., estimated *h*^2^_SNP_ *≈* 10%–30% while actual *h*^2^_SNP_ *≈* 70%–90%, can shift the future focus from genetic research toward environmental factors (*e.g*., diet, smoking, humidity, and temperature) and vice versa.

Phenotypic variance consists of two main components: genetic and environmental factors. Environmental physiochemical interaction with organisms and their influences on traits are well addressed aspects^3–5^. When phenotypic variation is observed for genotypes (individuals) within a single, specific environment, the study is considered a single-environment study. However, if observations are made across multiple environments, the study is classified as a multi-environment study, which is most relevant to animal and plant genetic research. In contrast, this concept does not apply to humans due to the lack of genotypic replication and controlled lifestyle or living conditions; individuals have their own unique lifestyle and environment. Similar to the inclusion of new genotypes, incorporating additional environments into a multi-environment study can alter the variance and heritability levels. Consistent with this, heritability shifts have been observed across different experimental conditions such as age^6^. In single-environment experiments, genetic variance is expected to account for the majority of phenotypic variance, approaching 100% heritability, with any deviations typically attributed to sampling errors. However, SNP heritability estimates in single-environment experiments are exceedingly underestimated. Experimental challenges that can affect heritability estimates include inadequate genotypic coverage, partial genomic coverage, imprecise phenotypic scoring scales, assignment of unassociated SNPs to estimation models, exclusion of associated SNPs from estimation models, neglecting epistatic interactions, and ignoring non-additive allelic effects. For example, approximately 3^100,000^ individuals with different genotypes are required for precise genetic estimations in a diploid genome with 100,000 biallelic SNPs, given that mutations are independent and conditional lethal SNPs are exceedingly rare. Various models have been developed to estimate SNP heritability, for example, using the minor allele frequency (MAF) and the linkage disequilibrium based weighted SNP effects^2,7–9^. Despite genotyping by millions of variants, SNP heritability still captures 30%–50% less phenotypic variance compared to heritability estimates derived from family and twin studies^1,9–19^. This underestimation is even more evident when compared to the much higher levels of heritability estimates, typically expected in single-environment studies that utilize millions of SNPs. Approaching total genomic and genotypic coverage, along with more robust scoring systems, such as subclasses for qualitative traits, can help bridge the gap. Capturing epistatic variance^20^ and controlling population structure^21^ have also been introduced to reduce the missing heritability levels. However, these strategies alone may not sufficiently close the gap unless SNPs are properly allocated to estimation models that account for both additive and non-additive effects between reference (*Ref*) and alternative (*Alt*) alleles.

Polymorphic genomic positions, should either be neutral (such as unassociated SNPs) or contribute positively (such as positive and negative associated SNPs) when included in models estimating SNP heritability. However, this study shed light on the negative contributions to heritability estimates imposed by SNPs with large association *p*-values through the commonly used genome-wide relatedness matrices. Family-based heritability estimations employ the true population statistic of genetic similarities (Fig. 1), *e.g*., monozygotic twins are genetically identical by descent and fraternal twins share, on average, 50% of their genomes^22^. In contrast, SNP heritability calculations rely on genome-wide estimates of genetic similarities, which are not phenotype-specific and are therefore susceptible to substantial error, as each phenotype is governed by a small fraction of the genome (Fig. 1). In Figure 1, we present an example where the addition of one unassociated SNP increases the genetic similarity from 50% (obtained based on an associated SNP) to 62.5%, introducing a downward bias in the phenotype-specific genetic variance. Despite the identification of this common miscalculation over a decade ago^23,24^, the majority of studies have continued to rely on the relatedness matrices derived genome-wide using all SNPs. It is evident that unassociated SNPs, *i.e*., SNPs from the upper-bound of a dataset specific *p*-value threshold, result in overestimation of genetic similarities (Fig. 1), ultimately leading to the underestimation of SNP heritability. Consistent with this, studies report remarkably low SNP heritability estimates (Average = 9.6%, Median = 6.6%, Fig. 2a). Accepting these estimates would confirm that many of the traits studied are purely environmental characteristics rather than genuine traits governed by genomes. This, of course, cannot be true and clearly indicates a downward bias in genetic variance estimates. Relying on a predefined *p*-values threshold such as 10^-8^ is also prone to underestimations of heritability given that polygenicity is a common phenomenon and many relevant variants may fall above this arbitrary cutoff. These technical challenges have led to the widespread practice of using either all SNPs or only those with *p*-value *≤* 10^-8^, depending on which approach yields the highest heritability estimate in every study. One of the remarkable examples is the heritability estimates for human height^15,16^. Furthermore, false-positive SNP associations and the neglect of non-additive genetic effects can lead to over-estimations and underestimations of heritability, respectively. This study is accordingly focused on excluding unassociated SNPs with negative contributions to heritability, minimizing false-positive SNP hits, and finally capturing both additive and non-additive effects between *Ref* and *Alt* alleles. We introduce SAFE-*h*^2^ (SNP Allocation For Estimating Heritability) method and application tool (Supplementary Fig. 1) that performs a *p*-value based SNP and heritability profiling to identify a dataset-specific *p*-values threshold, preventing biologically meaningless negative contributions to SNP heritability estimates. SAFE-*h*^2^ also defines a safe *p*-value interval to mitigate the heritability contributions of false-positive SNP hits. Furthermore, SAFE-*h*^2^ introduces an allelic adjustment strategy based on the observed genetic effects for every SNP to jointly capture intra-locus non-additive and additive allelic effects (*Ref* -allele_SNPi_ × *Ref* -allele_SNPi_, *Ref* -allele_SNPi_ × *Alt* -allele_SNPi_, and *Alt* -allele_SNPi_ × *Alt* -allele_SNPi_), enhancing the precision of SNP heritability estimates. SAFE-*h*^2^ can leverage heritability estimation models used in EMMAX^25^, LDAK-GCTA, LDAK-Thin^7^, GCTA-GREML^26^, and GEMMA^27^.

**Figure 1:**
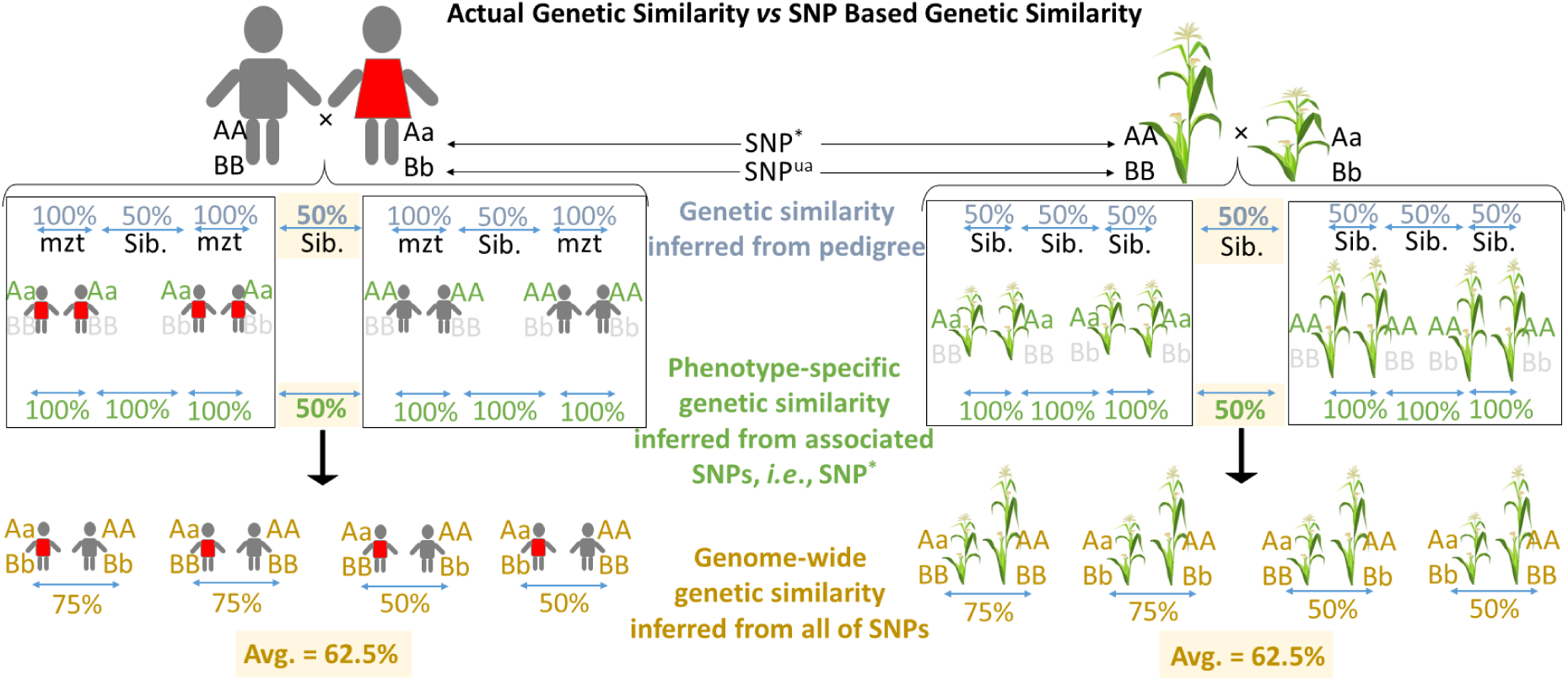
Unassociated SNPs contribute to a downward bias in genetic variance estimates. Among genotypes which are differentiated based on a given phenotype-of interest, family-based studies apply actual genetic similarities which are directly reflected in the genetic similarities inferred from associated SNPs (SNP^*^). However, genome-wide genetic similarities are overestimated when they are based on all SNPs, *i.e*., associated (SNP^*^) and unassociated (SNP^ua^). The mzt and Sib refer to monozygotic twins and siblings, respectively.

**Figure 2:**
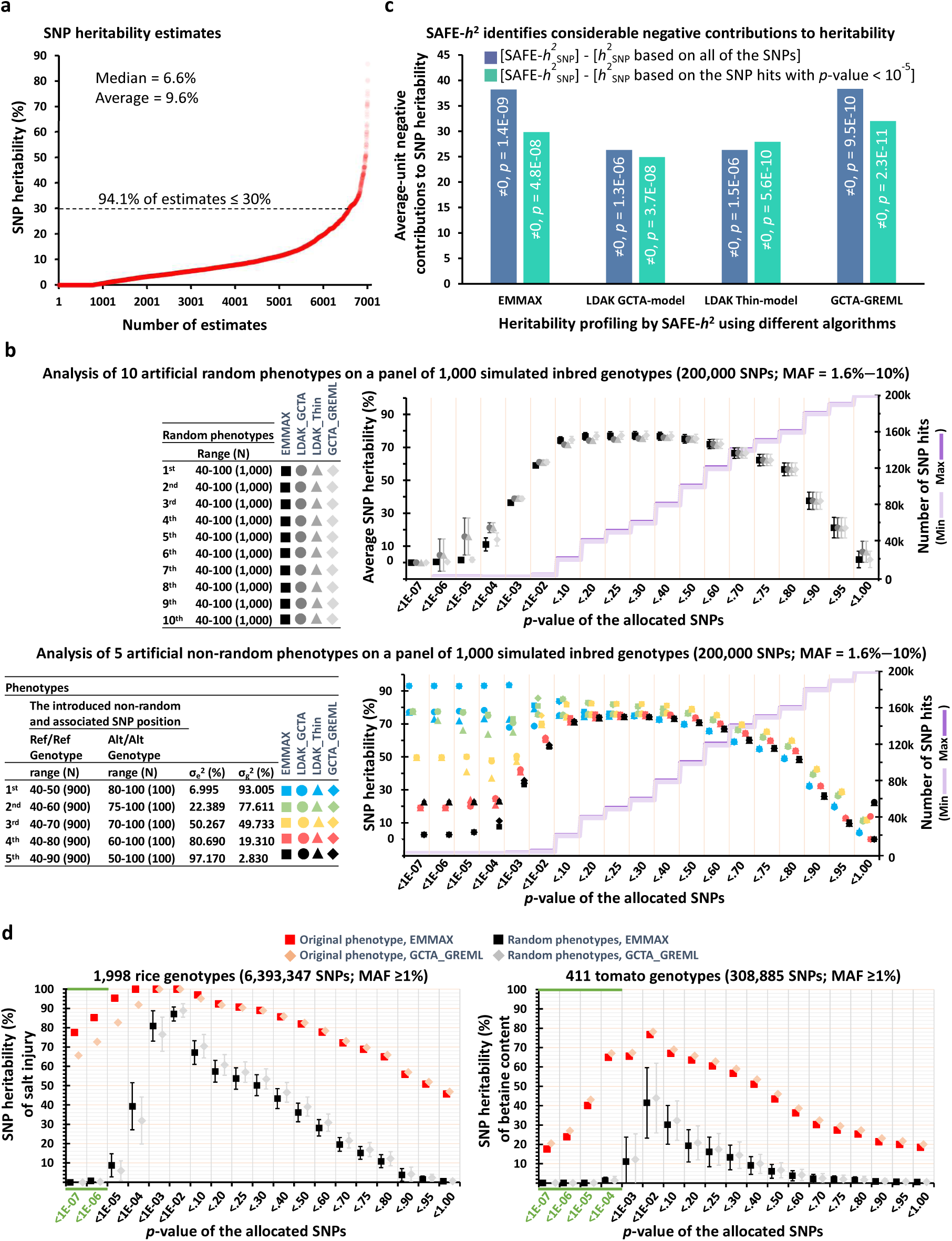
SAFE-*h*^2^ ensures accurate SNP heritability estimates by discarding meaningless negative contributions and restricting false-positive inputs. a) SNP heritability estimates were obtained from 110 recently published studies (see Supplementary Table 1), encompassing 7,008 independent estimates across various traits. b) EMMAX, LDAK-GCTA, LDAK-Thin, and GCTA-GREML were applied to estimate SNP heritability for the clusters of SNPs identified by their association *p*-values. By ensuring additive-only effects, five different levels of associations were simulated for a predesignated SNP position and as a result, the SNP heritability levels were predictable at or above the predefined levels, given that other SNPs could also, by chance, account for additional phenotypic variance. N represents the number of genotypes. The error bars represent 99% confidence intervals. c) SAFE-*h*^2^ improves the heritability estimations for 14 and 10 phenotypes in *Arabidopsis* and rice, respectively, by preventing the negative contributions of weakly-associated SNPs. The *p*-values are from two-tailed paired t-tests. d) SAFE-*h*^2^ minimizes false-positive contributions. The safe-intervals for *p*-value are highlighted in green representing the area with minimum false-positive-SNPs contribution to heritability. The average and standard deviation of estimations are illustrated for six random phenotypes simulated within the range of original phenotype. b–d) GCTA-GREML was performed based on the “inbred” algorithm. MAF: minor allele frequency, σ_g_^2^: inter-genotype variances, σ_e_^2^: intra-genotype variations.

## Results

### Within an array of association p-values, SNPs with large p-values neutralize the contribution of SNPs with small p-values to heritability estimates

The meaningless negative contributions to heritability depend on levels of association *p*-values in relation to each other. Therefore, we simulated phenotypes as random integers to generate arrays of association *p*-values, rather than representing true associations, for illustration of negative contributions. Analyses were performed on 1,000 and 10,000 simulated genotypes with 100, 200k, and 2,000k SNPs (MAF = 1.6%–10%) in linkage equilibrium as well as the human HapMap phase 1 genomic data^28^ with 1,092 individuals and 177,675 SNPs (MAF *≥* 1%) on chromosome 22. Analyses *via* four different models of EMMAX^25^, LDAK-GCTA, LDAK-Thin^7^, and GCTA-GREML^26^ revealed a consistent pattern of negative contribution to heritability estimates imposed by the SNPs having large association *p*-values (Fig. 2b, Supplementary Fig. 2a,c and 3). These analyses were replicated after incorporating 1, 12, 24, or 48 predesignated SNPs, each explaining 2.8% to 94.3% of the phenotypic variance through additive effects (Fig. 2b, Supplementary Fig. 2b,d and 4). It is virtually impossible to find multiple SNPs in linkage equilibrium and with large contributions to phenotypic variance and *p* values < 10^-20^ in real-world experiments with 200k SNPs and 1,000 genotypes. Nevertheless, there were 400 to 39,000 other SNPs with random associations and positive contributions to the heritability estimates among the 200k examined SNPs. Biologically implausible negative contributions were again observed for SNPs with large association *p*-values, leading to substantial underestimations of SNP heritability across all scenarios (Fig. 2b, Supplementary Fig. 2b,d and 4).

We observed the same pattern of negative contributions to heritability in linkage disequilibrium simulations, *i.e*., 200k SNPs on a 200kb chromosome (MAF = 1.6%–10%, Supplementary Fig. 5). Although, HapMap SNPs had MAFs up to 50%, we further simulated genotypes with MAFs ranging from 1.6%–50% and found the same patterns of negative contributions (Supplementary Fig. 6). Notably, LDAK-GCTA and LDAK-Thin models failed to accurately estimate the predefined heritability levels of 93.4%, 93%, 3.7%, and 2.8% when the estimations were solely based on the predesignated SNPs (*i.e*., *p* values < 10^-7^, Fig. 2b, Supplementary Fig. 2b,d and 4–6). In contrast, EMMAX and GCTA provided more stable estimates, yielding the closest approximations.

In any conceivable scientific experiment, replication serves to reaffirm the accuracy of the estimated statistics. When this confirmation cannot be achieved, it suggests an ineffective and flawed sampling/replication strategy. In other words, it indicates that the samples do not accurately represent the population from which they were drawn, as seen when comparing two datasets, both simulated independently from scratch. Therefore, it is important to note that performing replicated analysis—i.e., obtaining *p*-values from one simulated dataset and applying them to a second simulated dataset—is not a valid approach (see Supplementary Text 1). This is because adding a specific level of noise to the second dataset will result in the same estimates, plus or minus the introduced noise, which does not provide any additional information beyond what is already presented in Figure 2b and Supplementary Figures 2-6. Furthermore, *p*-values from one simulated dataset cannot be applied to another dataset that is independently simulated from scratch, as this would involve using *p*-values derived for one phenotype (e.g., weight) to estimate heritability for a different phenotype (e.g., a specific disease).

### SAFE-h^2^ effectively prevents the illogical negative contributions of SNPs to heritability estimations

To ensure accurate estimation of SNP heritability, we developed SAFE-*h*^2^ to perform SNP heritability profiling across *p*-value-guided SNP clusters, identifying an optimal *p*-value threshold for excluding unassociated SNPs with negative contributions. As a model plant, different traits have been studied through GWAS in *Arabidopsis*^3,29^. Here, heritability profiling was conducted for 14 phenotypes in *Arabidopsis* using 2,621k to 2,840k SNPs^30–32^ as well as 10 phenotypes in rice using 6,131k to 6,414k SNPs^33–35^. SAFE-*h*^2^ improved the accuracy of heritability estimates by excluding negative contributions. Notably, SNPs with negative contributions masked the heritability estimates by an average of 32.3 units (5.6–64.7, Fig. 2c, Supplementary Table 2). Heritability levels were also underestimated by an average of 28.7 units (13.6–63.7) when based solely on SNPs with *p*-values *≤* 10^-5^ (Fig. 2c, Supplementary Table 2). We used SAFE-*h*^2^ to analyze 50 additional phenotypes from three plant and three animal species^36–44^ and found SNPs with large association *p*-values imposing an average negative contribution of 30 units to the heritability estimates (Supplementary Table 3). SAFE-*h*^2^ also outperformed BayesR estimator^45^ in the simulated (linkage equilibrium and linkage disequilibrium scenarios) and the real datasets. BayesR either failed to detect or underestimated the genetic variances, while being 8–10 times slower (Supplementary Tables 4–6).

### SAFE-h2 reports a safe p-value interval to minimize positive contributions of false-positive SNP hits to heritability estimates

False-positives are particularly challenging not only because of the resource-intensive nature of GWAS but also due to the impracticality of near-complete (*≈*2^(no. SNPs)^ or *≈*3^(no. SNPs)^ in the haploid or the diploid state, under a biallelic scenario) genotypic coverage. SAFE-*h*^2^ uses simulated phenotypes to pinpoint a safe *p*-value interval with minimum SNP heritability level at the lower-tail of heritability profile. For example, we can set the *p*-value interval from 0 up to a maximum *p*-value, determined at a user selected noise level, e.g., 0% or 5%. The heritability noise levels are obtained by analyzing simulated phenotypes within the range of the phenotype-of-interest. The SNP heritability for the original phenotype is subsequently estimated at the upper-end of this interval to minimize false-positive contributions. Heritability estimates using all SNPs were consistently underestimated not only when compared to the SAFE-*h*^2^ heritability estimates with excluded negative contributions but also when compared to the SAFE-*h*^2^ heritability estimates with excluded negative and false-positive contributions (Fig. 2d, Supplementary Fig. 7).

### SNP heritability estimation through joint-analysis of intra-locus allelic effects of additive and non-additive

The significance of intra-locus non-additive allelic effects (*Ref* -allele_SNPi_ × *Ref* -allele_SNPi_, *Ref* -allele_SNPi_ × *Alt* -allele_SNPi_, and *Alt* -allele_SNPi_ × *Alt* -allele_SNPi_) has been reviewed^46^ and methods like GCTA-GREMLd has been introduced to address dominance effects in heritability estimations^47^. However, the estimated small intra-locus non-additive effects have led to the widespread assumption that these effects are negligible^46^. Similar to pervasive non-linear interaction of genes^20,48^, there is no inherent genetic or biological rationale and the estimated small non-additive effects between reference and alternative alleles may be attributable to methodological limitations and artifacts, aligning with our results.

When linear models are augmented with some allelic adjustments tailored to the observed non-additive effects, they capture genetic variance at levels comparable to those of the exponential model (Fig. 3a–f). SAFE-*h*^2^ employs the same allelic adjustment algorithm to extract non-additive effects within the framework of linear models. For every SNP position, SAFE-*h*^2^ examines different scenarios, *i.e.*, additive (original genotypes), adjusted genotypes for dominance effects, adjusted genotypes for overdominance effects, and adjusted genotypes for heterosis-like effects, to find the best fitted linear model. Before heritability profiling, combinatory genotype datasets can then be built based on one selected adjustment or the original genotype, *i.e*., the one with smallest *p*-value, for every SNP position. The SAFE-*h*^2^ allelic adjustment improved the SNP heritability estimations up to 9.7 units for the 10 random phenotypes assigned to 1,092 HapMap genotypes (Fig. 3g), as well as for 10,000 simulated genotypes derived from the 1,092 HapMap genotypes (Fig. 3h). In contrast, the GCTA-GREMLd either showed no improvement or resulted in an average underestimation of 3.46 units compared to the additive approach (Fig. 3g,h). The SAFE-*h*^2^ allelic adjustment proved robust enough that no further improvements in heritability estimates were observed even after increasing the cohort size from 1,092 to 10,000 (Fig. 3i). On the contrary, the additive and GREMLd based estimates were improved after cohort enlargement (Fig. 3i). The SAFE-*h*^2^ additive and non-additive joint-analysis was also evaluated using a biallelic SNP associated at different rates and through additive, dominance, overdominance, and heterosis-like genetic effects to simulated phenotypes (Table 1). When the underlying genetic effects were additive, the estimates of SNP heritability based on joint additive and non-additive effects were identical to those derived solely from additive effects (Table 1). When the underlying genetic effects were non-additive, the allelic adjustments for non-additive effects by SAFE-*h*^2^ improved the heritability estimations using all the four models, *i.e.*, EMMAX, GCTA-GREML, LDAK-Thin, and LDAK-GCTA (Table 1). GCTA-GREML and LDAK were however less effective at capturing additive-only or joint additive and non-additive effects at different association levels (Table 1). SAFE-*h*^2^ obtained the best approximates of the genetic variances, *i.e.*, here equivalent to the broad-sense heritability, when applying the EMMAX and GEMMA models (Table 1, Supplementary Table 7). SAFE-*h*^2^ improved estimates by capturing 6.22% extra broad-sense heritability on the allelic effects of dominance, 16.22% extra broad-sense heritability on the allelic effects of overdominance, and 24.31% extra broad-sense heritability on the heterosis-like effects on average. Analyzing 50 phenotypes observed in plant and animal species^36–44^, further validated the SAFE-*h*^2^ allelic adjustment by extracting non-additive effects as well as the SAFE-*h*^2^ superior and stable performance versus GREMLd (Fig. 3j, Supplementary Fig. 8). It is important to emphasize that the overestimation of SNP heritability can only occur through positive contributions from false-positive SNP hits, which depends on statistical power levels. At an identical statistical power level, *i.e.*, identical genomic and genotypic coverage, the most effective heritability estimation method is the one that captures the greatest proportion of phenotypic variation.

**Figure 3:**
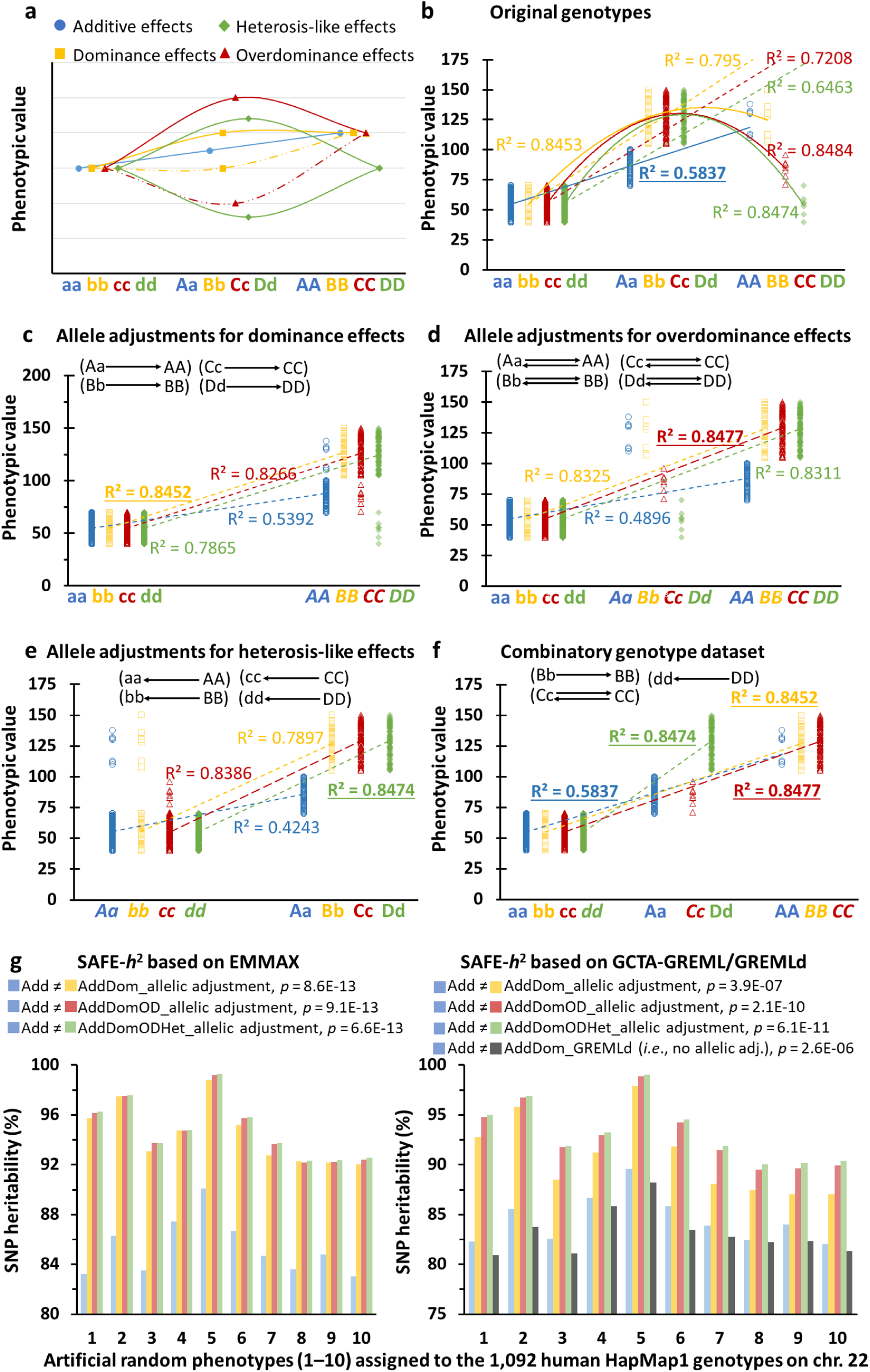
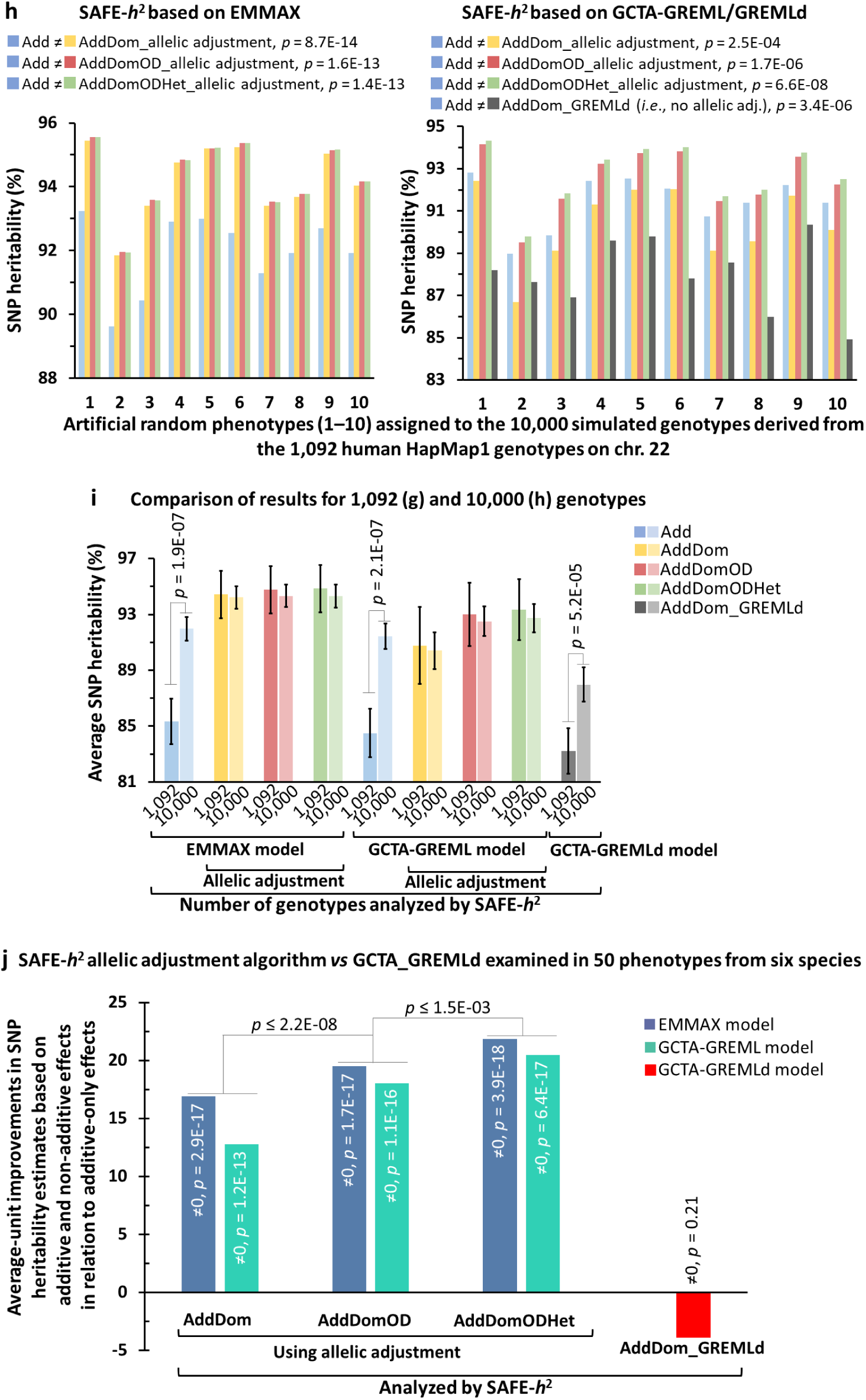
Allele adjustments capture intra-locus non-additive allelic effects. a–f) Demonstration of four biallelic loci; locus A/a with additive effect, locus B/b with dominance effect, locus C/c with overdominance effect, and locus D/d with heterosis-like effect. The best linear-fits are highlighted by bold underlined R^2^ values. The adjusted alleles are in italic. Single arrows indicate allele adjustment by conversion and double arrows indicate allele adjustment by replacement. a) Allelic effects in a biallelic scenario. Dominance and overdominance effects can be on either alternative or reference alleles (*Alt* or *Ref*) which are differentiated by continues and dashed extrapolations. b) The artificial phenotype no. 3 (see Supplementary Fig. 2d), and its derivatives were used to represent the additive and non-additive effects, respectively. Continues lines represent the most fitted models; this was linear for the locus A/a. Low-fit linear approximates (dashed lines for B/b, C/c, and D/d loci) are also shown. c) Allelic adjustment for the dominance effects of Ref alleles (A, B, C, & D) leads to an improved linear-fit only on the B/a locus having dominance effect. d) Allelic adjustment for the overdominance effects of Ref alleles (A, B, C, & D) leads to an improved linear-fit only on the C/c locus having overdominance effect. e) Allelic adjustment for the heterosis-like effects leads to an improved linear-fit on the D/d locus with heterosis-like effect. f) A combinatory genotype dataset contains every SNP position in either of its original or one of the adjusted states, depending on which provides the best linear-fit, *i.e.*, highest R^2^. g–i) Capturing additive and non-additive effects for the 10 artificial random phenotypes described in Supplementary Fig. 2c. j) The SAFE-*h*^2^ allelic adjustment algorithm efficiently captures additive and non-addtive effects. g–j) The *p*-values are from two-tailed paired t-tests. The error bars represent 95% confidence intervals. Add: heritability based on additive effects, AddDom: heritability based on additive and dominance effects, AddDomOD: heritability based on additive, dominance and overdominance effects, AddDomODHet: heritability based on additive, dominance, overdominance, and heterosis-like effects, GREMLd: heritability estimated by capturing additive and dominance effects without allelic adjustment, MAF: minor allele frequency.

**Table 1:**
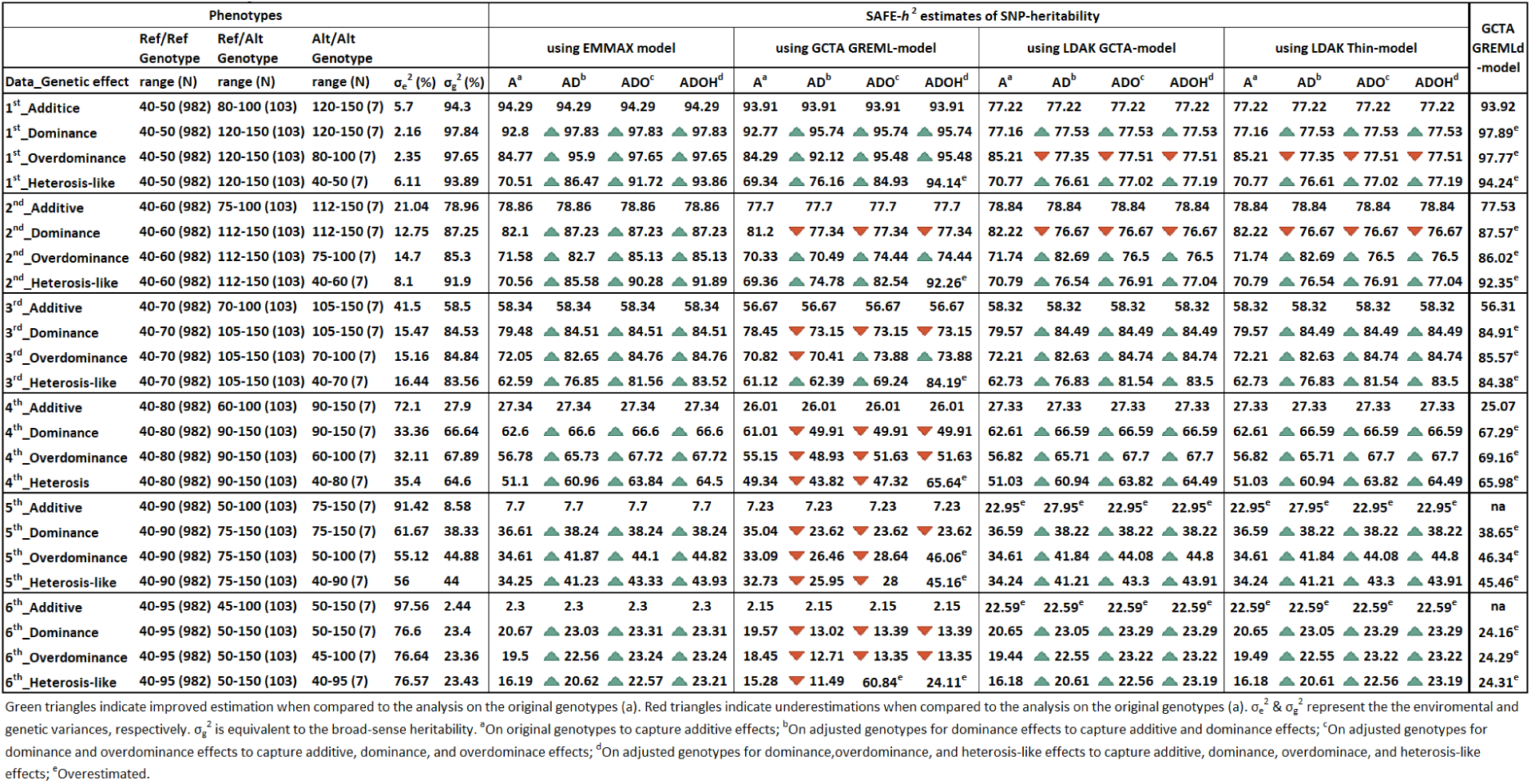
SNP heritability estimations on additive or additive + non-additive effects of a biallelic locus.

## Discussion

SNP heritability estimations are typically obtained by including all SNPs or by selecting SNPs with *p*-value less than 10^-8^, often depending on which approach yields the highest estimate. A remarkable example is the heritability estimates on human height^15,16^. Functional, regional, linkage disequilibrium, and MAF stratified SNPs have also been employed in SNP heritability calculations^8–10,12,13,15^. Nevertheless, these approaches provide estimates of genetic variance at sub-genome levels rather than phenotype-specific estimates (*i.e.*, associated SNP-specific estimates). It is evident that all of these SNP heritability estimation strategies are vulnerable to the inclusion of unassociated SNPs with upper-tail association *p*-values, potentially introducing illogical negative contributions to heritability estimates. Although the genetic relatedness matrices ideally require phenotype-associated SNPs and not irrelevant variants, the trade-off between the detrimental effects of excluding associated SNPs and including irrelevant variants has made the SNP selection challenging, often resulting in inaccurate genomic predictions, also addressed previously by Lippert et al. (2013)^24^. The SAFE-*h*^2^ method bypasses this trade-off by identifying a *p*-value threshold below which no SNP can be found with deflationary effects on genetic variance. Finally, SAFE-*h*^2^ generates a list of *p*-value-guided SNP clusters and outputs a profiling graph of heritability estimates (Supplementary Fig. 9a–c). The EMMAX and GEMMA algorithms were found as the most stable models for accurate estimation of SNP heritability using SAFE-*h*^2^ (Fig. 2, Table 1, Supplementary Fig. 2,4–6, Supplementary Table 7). For EMMAX-based estimations, SAFE-*h*^2^ utilizes the Balding-Nichols (BN) kinship matrix, which has been shown to be highly correlated with FaST-LMM and GEMMA kinship matrices^49^. Compared to the analyses based on *p*-values from EMMAX, we observed even larger negative contributions to heritability estimations when the analyses were based on *p*-values calculated by GEMMA (Supplementary Table 8), which employs an exact mixed model^27^.

A second key challenge in GWAS studies is the low statistical power due to insufficient genotypic coverage, leading to false-positive results. SAFE-*h*^2^ enables the identification of a second *p*-value threshold at which contributions of false-positive SNP hits to heritability estimations can be restricted efficiently (Fig. 2d, Supplementary Fig. 7 and 9e). This approach can also be applied in GWAS to define significant SNPs. The hidden heritability due to false-negative SNP hits has also been discussed previously^50^. SAFE-*h*^2^ ensures the inclusion of weakly associated SNPs, which might otherwise be excluded (*i.e*., leading to false-negatives) due to a predefined *p*-value cut-off, *e.g*., 10^-8^. Replication of experiments, *e.g.,* using effect-sizes of associated SNPs obtained in one cohort to calculate polygenic risk scores for individuals/genotypes in a second cohort, is a common approach in GWAS to mitigate overfitting^51^ caused by inadequate genotypic coverage. However, replication to reduce overfitting is not applicable in heritability estimations. It is important to note that overfitting occurs at genotype-level, while heritability is a population- or cohort-level statistic that is specific to datasets and experiments, varying with new genotypic and environmental inputs, as outlined in the introduction. An interesting example in this case is the observed heritability shifts across different experimental conditions such as age^6^.

The recently developed tools such as Next-Gen GWAS^20^ and GWAIS^52^ offer major improvements in the processing of epistatic (SNP × SNP) effects. To complement, SAFE-*h*^2^ is capable of extracting intra-locus (*i.e*., *Ref* -allele_SNPi_ × *Ref* -allele_SNPi_, *Ref* -allele_SNPi_ × *Alt* -allele_SNPi_, and *Alt* -allele_SNPi_ × *Alt* -allele_SNPi_) non-additive allelic effects. An example of SAFE-*h*^2^ output graphs for joint analysis of additive and non-additive allelic effects is shown in Supplementary Figure 9d. SAFE-*h*^2^ achieved the best performance using EMMAX and GEMMA models to jointly capture additive and non-additive allelic effects of the predefined single biallelic-locus, without overestimations (Table 1, Supplementary Table 7). In contrast, GCTA-GREMLd, while still providing useful estimates, resulted in small overestimation errors across all measurements (Table 1). GCTA-GREMLd was also prone to SNP heritability underestimations, as its estimates for joint additive and non-additive effects were smaller than those for additive-only effect (Fig. 3g,h,j, Supplementary Fig. 8). Furthermore, SAFE-*h*^2^ extracts not only the dominance but also overdominance and heterosis-like allelic effects.

In summary, SAFE-*h*^2^ provides phenotype-specific estimates of SNP heritability rather than genome or sub-genome specific estimates, and minimize the likelihood of false-positive contributions. SAFE-*h*^2^ further refines the estimations through joint analysis of intra-locus non-additive and additive allelic effects. By adjusting alleles, SAFE-*h*^2^ generates genotypic datasets that can facilitate linear association tests to address intra-locus additive and non-additive effects jointly.

## Materials and Methods

### Datasets

The simulated homozygote genotypes were arranged across 29 chromosomes, with 300kb distance between SNPs to meet linkage equilibrium. For each genotype, reference and alternative alleles were randomly assigned at different chromosomal positions. VCF files for 1,000 and 10,000 genotypes were assembled with 200k SNPs with MAF ranging from 1% to 10%. Within these datasets, one of the 200k SNP positions was designated as non-random to be associated with the simulated non-random phenotypes. For having multiple predefined associations, 11, 23, or 47 extra predefined SNPs were integrated into the same genomic datasets. Genotypes with 100 and 2,000k SNPs were also simulated. Further proof-of-concept analyses were accomplished using human chromosome 22 from the HapMap phase 1 genomic data^28^. This included 1,092 genotypes and 177,675 SNPs after dataset filtering (MAF *≥* 1%). As for the simulated genotypes, one simulated non-random SNP position was introduced among the HapMap variants to be associated with the simulated non-random phenotypes. The human chr.22 SNP-set was also extended by simulation to 10,000 individuals. For this, the original SNP dataset was shuffled three times among the chromosomal positions to get a new genotype set. This was repeated to have 9 new datasets. The original and new datasets were then horizontally assembled to build a dataset of 10,000 genotypes represented by 177,675 SNPs. The final simulated dataset had 177,445 variants after filtering for MAF of *≥* 1%. The artificial phenotypes were produced as random or non-random integers assigned randomly among the genotypes or among the genotypic groups of the simulated SNPs with varying levels of associations. Comprehensive analyses were also conducted using *Arabidopsis* and rice public datasets. For *Arabidopsis*, 14 different phenotypes were obtained from the AraPheno database^30^. These phenotypes were from a panel of 328 to 1,058 genotypes. The imputed genomic variants for these panels^31,32^ contained 2,621,362 to 2,840,072 post-filtering SNPs. The public rice data was obtained from the rice SNP-Seek database^33^. The analyses were performed using 10 phenotypes that were previously recorded in 1,422 to 2,102 rice genotypes. The rice genomic data was also publicly available in rice SNP-Seek database^33–35^. The genotypes contained from 6,131,325 to 6,414,800 post-filtering SNPs. Additional datasets from both animal and plant species were analyzed to examine the negative contributions to heritability as well as the inclusion of intra-locus allelic effects of non-additive in the SNP heritability estimates. These included clutch size in 314 wild avians^36^ genotyped by 40,766 SNPs (MAF *≥* 0.01), four fruit and flowering related phenotypes in 96 tomato genotypes^37^ genotyped by 16,782 SNPs (MAF *≥* 0.01), the contents of 15 secondary metabolites in 411 tomato genotypes^43^ genotyped by 308,885 SNPs (MAF *≥* 0.01), the contents of 7 secondary metabolites in 136 apple genotypes^38^ genotyped by 98,584 SNPs (MAF *≥* 0.01), apple fruit-weight, apple sucrose, fructose, and tartrate contents in 368 to 461 genotypes^39^ represented by 3,055,912 SNPs (MAF *≥* 0.01), six growth related phenotypes in 752–755 goat genotypes^40^ represented by 40,639 SNPs (MAF *≥* 0.01), the abundance of 12 microbiome sub-communities in 827 genotypes of foxtail millet represented by 161,562 SNPs^44^, and a merged dataset for susceptibility to diabetes mellitus in 1,022 genotypes represented by 826 SNPs (MAF *≥* 0.01) that derived from two independent studies in cats^41,42^.

### Data analysis

Plink^53^ and EMMAX^25^ were used for the initial processing of genomic data and association analyses. MAF was *≥* 1% in all the analyses. A complete record on the latitudes of geographical origins was available for *Arabidopsis* genotypes when analyzing the stem length and leaf mineral contents. This geographical information was applied as quantitative covariates in the analyses. SAFE-*h*^2^ was developed to perform a dynamic SNP allocation for estimation of narrow-sense heritability. A comprehensive manual is provided together with the software. In brief, SAFE-*h*^2^ begins by scanning the input files and executing the user-defined combination of heritability estimation models on *p*-value guided SNP clusters. SAFE-*h*^2^ identifies a dataset- and experiment-specific *p*-value threshold at which SNPs with larger *p*-values are distributed randomly across genotypes in relation to the phenotypic variation, imposing negative contributions to heritability estimates. In essence, this *p*-value threshold is the best border that we can assume between associated and unassociated SNPs under standard experimental conditions, *i.e.*, the genomic and genotypic coverage, for accurate estimation of SNP heritability. Therefore, we define these SNPs with *p*-values larger than the threshold as “unassociated”. These unassociated SNPs lead to overestimated genetic similarities within genetic relationship matrices and thereby, underestimations in genetic variance and SNP heritability levels. SAFE-*h*^2^ requires a *p*-value file for SNPs, the Plink b-files (“bim”, “bed”, and “fam”), and optional covariate files specific for every estimation model. SAFE-*h*^2^ leverages five different models of EMMAX^25^, LDAK-GCTA, LDAK-Thin^7^, GCTA-GREML^26^, and GEMMA^27^ to estimate SNP heritability. GCTA-GREML inbred algorithm can also be applied to inbred genotypes or populations with considerable levels of inbreeding. To operate, SAFE-*h*^2^ requires the binary files of the mentioned tools (see the SAFE-*h*^2^ manual with full description). In return, SAFE-*h*^2^ generates the output data files and related graphs for clustered SNP hits and heritability profiles. It also features the *p*-value threshold and the estimated SNP heritability at that threshold on the graphs. Given that different levels of false-positive rates are typically expected in GWAS, SAFE-*h*^2^ employs an innovative approach to exclude these false-positive SNP hits. SAFE-*h*^2^ determines a “safe” *p*-value interval, with its upper-end to be used as an alternative *p*-value threshold for heritability estimations, ensuring minimal contributions from false-positive SNP hits. To perform the analysis, the user should simulate six independent random phenotypes within the range of the original phenotype-of-interest for all genotypes in the dataset. SAFE-*h*^2^ requires several input files: a “bim” file, a “bed” file, a “fam” file containing the original phenotypic records, six “fam” files for simulated random phenotypes, SNP *p*-value files for the real (one file) and random phenotypes (six files), and optional covariate files specific for estimation models. Using these input files, SAFE-*h*^2^ creates the SNP heritability profiles for all the random phenotypes as well as the real phenotype-of-interest using an identical genomic dataset. Users can select the estimation models of interest. Analyzing the random phenotypes subsequently determines a safe *p*-value interval with minimum SNP heritability level at the lower-tail of heritability profile. The upper-end of this interval can function as an alternative *p*-value threshold to omit the false-positive SNP hits from the estimation models and keep their contribution to heritability at minimum. Therefore, SAFE-*h*^2^ keeps false-negative and false-positive rates at minimum by heritability profiling for the phenotype-of-interest and the simulated phenotypes, respectively. Finally, SAFE-*h*^2^ generates an aligned graph of heritability profiles for the random and original phenotypes. The SNP heritability of the original phenotype can accordingly be defined at upper-end of the determined *p*-value interval to keep the false-positive contributions at its minimum.

SAFE-*h*^2^ also offers allelic adjustments for linear approximation on intra-locus allelic effects including dominance, overdominance, and heterosis-like effects. This facilitates capturing of non-additive allelic effects. SAFE-*h*^2^-preADOH builds two VCF files for the dominance adjustments by converting the heterozygote nucleotide positions into one of the homozygotes. It also creates two VCF files for the overdominance adjustments, where heterozygote nucleotide positions are converted into either of homozygotes and, the same homozygotes are converted into their heterozygote counterparts. For the heterosis-like effects, SAFE-*h*^2^-preADOH builds an additional VCF file by converting *Ref* × *Ref* homozygotes into the corresponding homozygote of *Alt* × *Alt*, and vice versa, at each nucleotide position. Here, the user can perform an independent association test or alternatively can command SAFE-*h*^2^-preADOH to execute the Plink’s glm function on the five adjusted and one original genotype datasets. To construct the combinatory genotype datasets, SAFE-*h*^2^-preADOH applies the association *p*-values to select the best linear-fit (with smallest association *p*-value) among the original and the five adjusted versions for every SNP position. At the end, SAFE-*h*^2^-preADOH provides three genotypic datasets that can be used to estimate the additive-dominance effects, additive-dominance-overdominance effects, and additive-dominance-overdominance-heterosis like effects. Here, every SNP-ID gets either of the following specific tags; [Add] for having additive effects on the phenotype of interest, [Dom] or [dom] for having dominance effects on the phenotype of interest, [OD] or [od] for having overdominance effects on the phenotype of interest, or [Het] for having heterosis-like effects on the phenotype of interest. Finally, SAFE-*h*^2^-ADOH performs the heritability profiling as explained for SAFE-*h*^2^ and on the three adjusted genotype datasets as well as the original genotype dataset. It is important to emphasize that SAFE-*h*^2^ considers very SNP through either of additive, dominance, overdominance, or heterosis-like effects for inclusion in the heritability model. Therefore, the file for additive-dominance effects is used to examine the SNPs tagged with [Add] only for additive effect and the SNPs tagged with [Dom] or [dom] only for dominance effect. The file for additive-dominance-overdominance effects is used to examine the SNPs tagged with [Add] only for additive effect, the SNPs tagged with [Dom] or [dom] only for dominance effect, and the SNPs tagged with [OD] or [od] only for overdominance effect. Finally, the file for additive-dominance-overdominance-heterosis effects is used to examine the SNPs tagged with [Add] only for additive effect, the SNPs tagged with [Dom] or [dom] only for dominance effect, the SNPs tagged with [OD] or [od] only for overdominance effect, and the SNPs tagged with [Het] only for heterosis-like effect.

## Author contributions

B.D conceived the study. B.D performed the analyses and compiled the SAFE-*h*^2^ script. B.D and M.N discussed the experimental designs and results. B.D, and M.N wrote the manuscript.

## Competing interests

The authors declare no competing interests.

## Acknowledgments

We thank Prof. Detlef Weigel (professor in plant genetics and evolution, Department of Molecular Biology, Max Planck Institute for Biology Tübingen, Germany) for his invaluable consult and comments.

## Code availability

SAFE-*h*^2^, containing the compiled SAFE-*h*^2^ scripts with related installation and usage manual, is attached to this publication as suppl. File 1. It also contains example datasets. SAFE-*h*^2^ is available for download at https://github.com/SAFE-h2/SAFE-h2-v2024-1. The example dataset can also be obtained from figshare (DOI: 10.6084/m9.figshare.24025728, Link: https://figshare.com/s/7193d332894f11deb1a5).

## Supplementary figures, tables, and text

**Supplementary Fig. 1:**
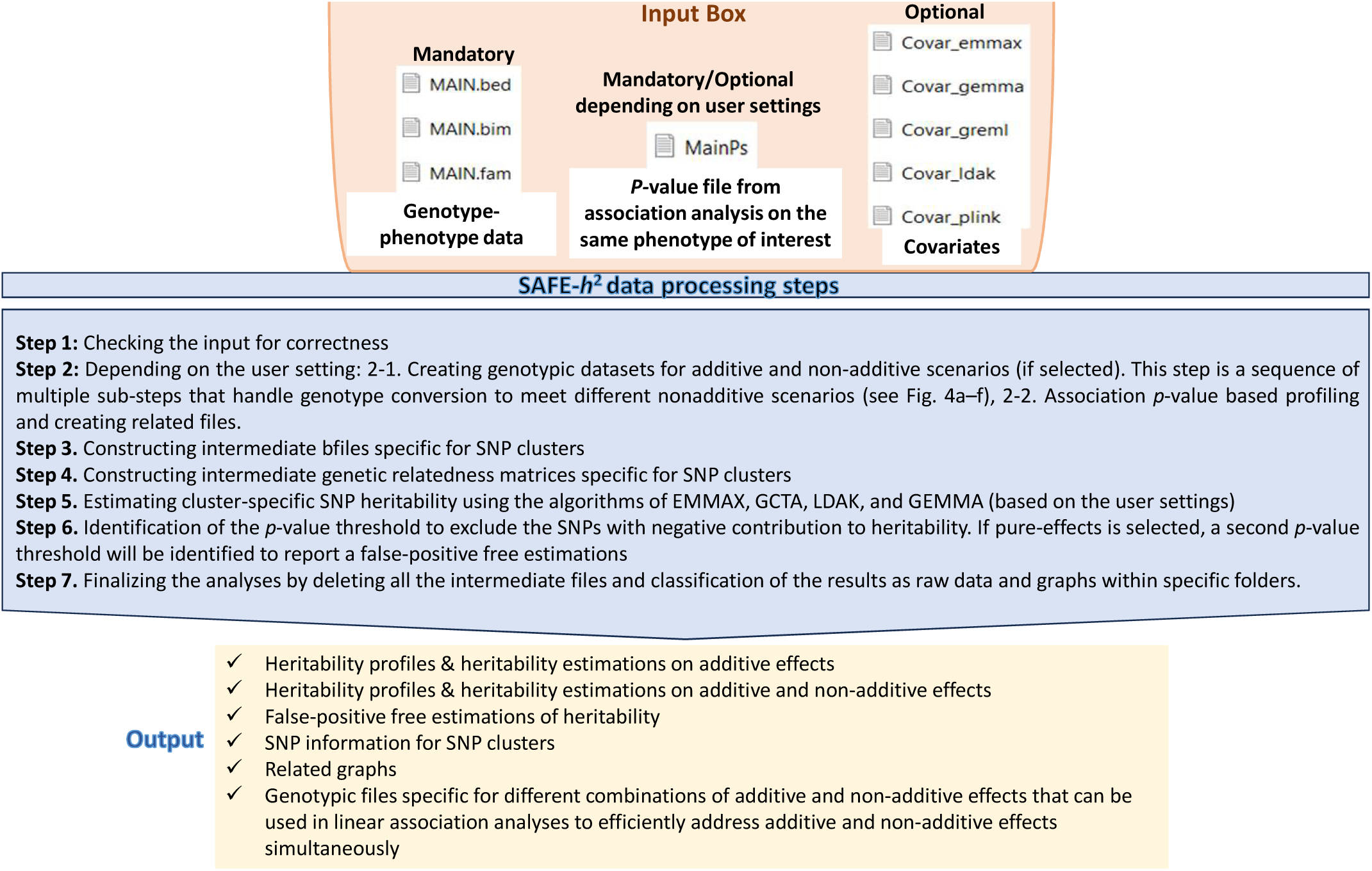
SAFE-*h*^2^ pipeline.

**Supplementary Fig. 2:**
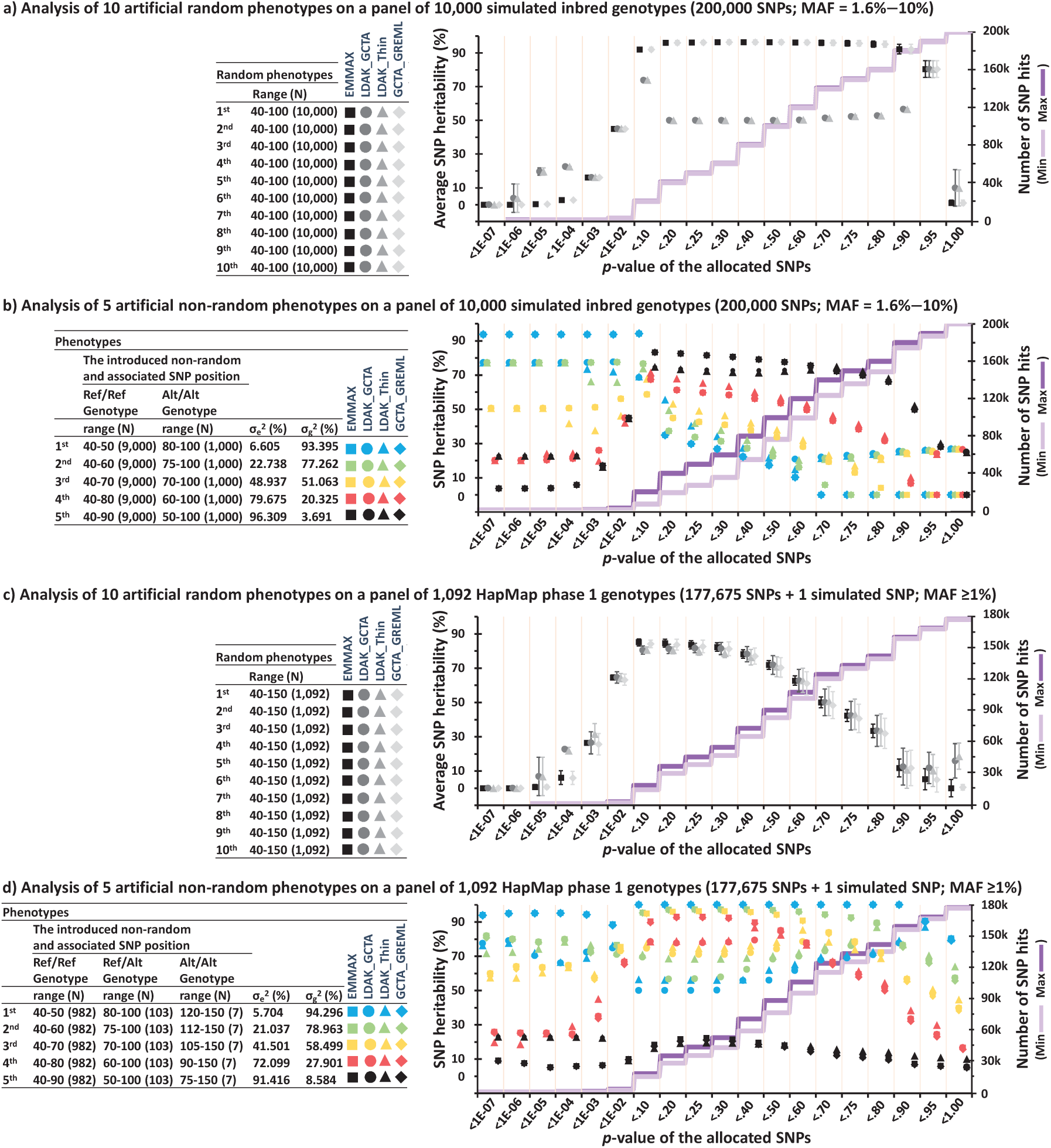
The SNP heritability profiles for simulated quantitative-phenotypes using simulated SNPs or variants of chr. 22 from the human HapMap phase 1 genotypes. a–d) An initial GWAS was performed using EMMAX to calculate association levels for the SNPs. The four different approaches of EMMAX, LDAK-GCTA, LDAK-Thin, and GCTA-GREML were then applied to estimate SNP heritability for the clusters of SNP hits identified using the association *p*-values. N represents the sample sizes, *i.e.*, number of genotypes. a,c) The average SNP heritability estimates were obtained for 10 independent random phenotypes within identical ranges. The error bars represent 99% confidence intervals. b,d) The SNP heritability estimates were obtained for five simulated phenotypes which were associated with a predesignated SNP position introduced within both the simulated and HapMap genomes. By ensuring additive-only effects, associations were simulated at five different levels. As a result, the SNP heritability levels were predictable at or above the predefined levels, given that other SNPs could also, by chance, account for additional phenotypic variance. a–d) For inbred genotypes, GCTA-GREML were performed based on the “inbred” algorithm. MAF: minor allele frequency, σ_g_^2^: inter-genotype variances, σ_e_^2^: intra-genotype variations.

**Supplementary Fig. 3:**
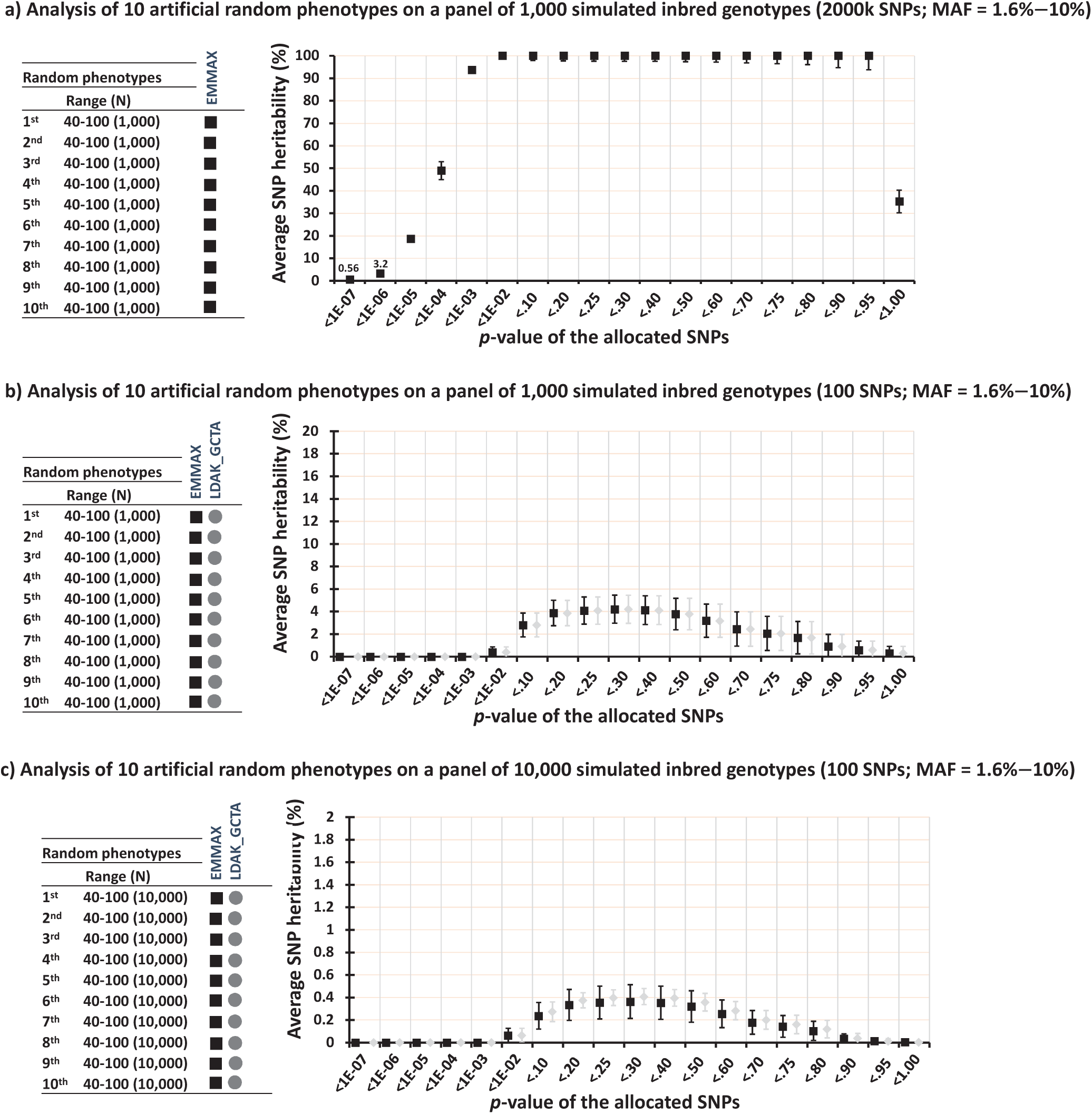
The SNP heritability profile for simulated random phenotypes (quantitative) and SNPs. An initial GWAS was performed using EMMAX to determine the association levels for the SNPs. The four different approaches of EMMAX, LDAK-GCTA, LDAK-Thin, and GCTA-GREML (by “inbred” algorithm) were then applied to estimate the SNP heritability for the clusters of SNP hits captured using the association *p*-values. N represents the sample sizes, *i.e.*, number of genotypes. MAF stands for the minor allele frequency. The average SNP heritability estimates were obtained for 10 independent random phenotypes within identical ranges. The error bars represent 99% confidence intervals.

**Supplementary Fig. 4:**
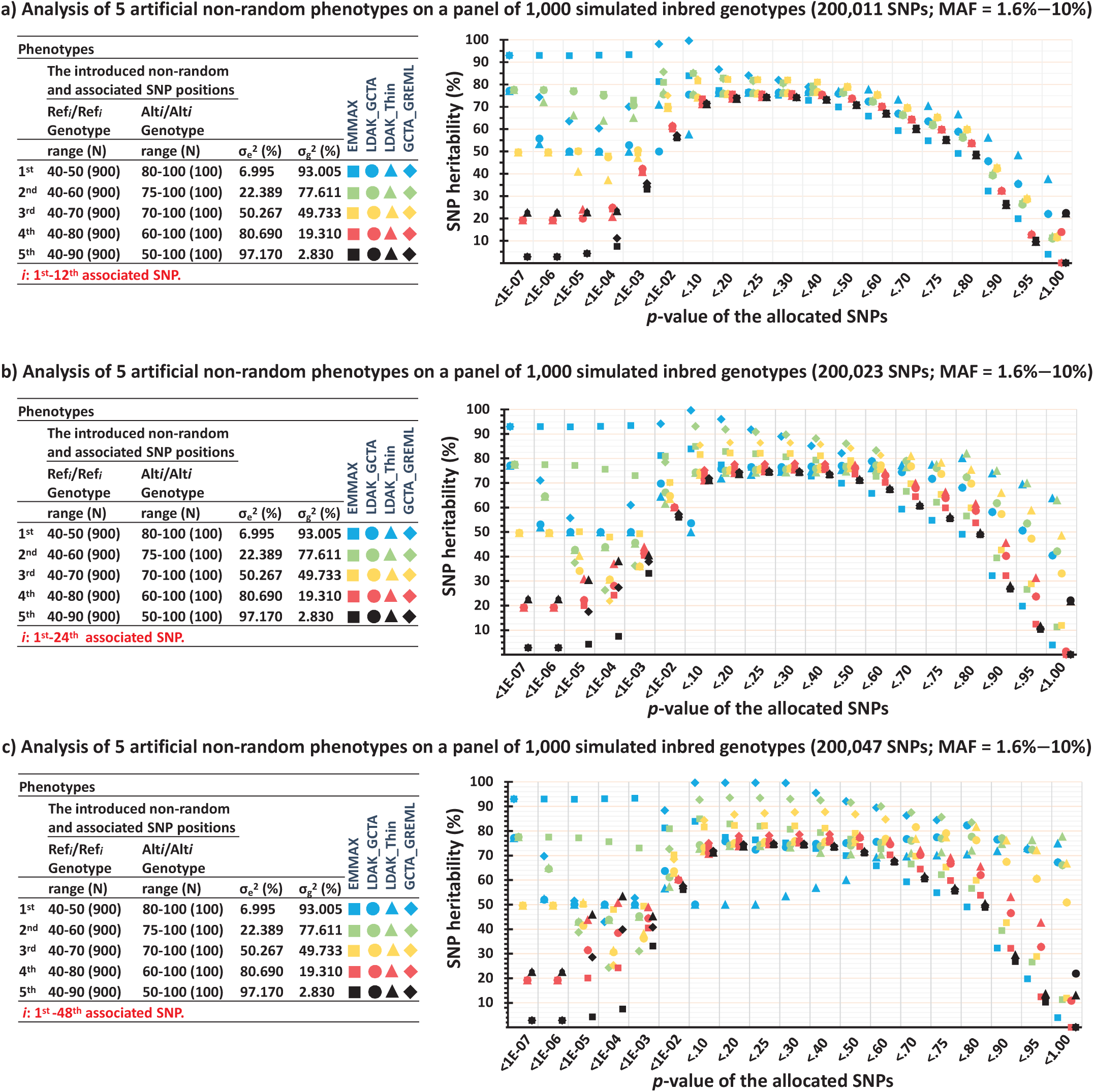
The SNP heritability profile for simulated quantitative-phenotypes and SNPs, with predesignated SNPs associated at varying rates. The SNP heritability estimations were predictable at or above the predefined levels, given that other SNPs could also, by chance, account for additional phenotypic variance. An initial GWAS was performed using EMMAX to determine the association levels for the SNPs. The four different approaches of EMMAX, LDAK-GCTA, LDAK-Thin, and GCTA-GREML (by “inbred” algorithm) were then applied to estimate the SNP heritability for the clusters of SNP hits captured using the association *p*-values. N represents the sample sizes, *i.e.*, number of genotypes. MAF stands for the minor allele frequency. Ensuring additive-only effects, the non-random phenotypes were defined among the genotypic groups of the introduced associated SNP positions, *i.e.*, Ref*i*/Ref*i* and Alt*i*/Alt*i* for the simulated inbred genotypes. The intra-genotype assignment of the phenotypes, while being within their specific ranges, was random. The inter-genotype variances of the phenotypes are shown as σ_g_^2^. σ_e_^2^ represents the intra-genotype variations.

**Supplementary Fig. 5:**
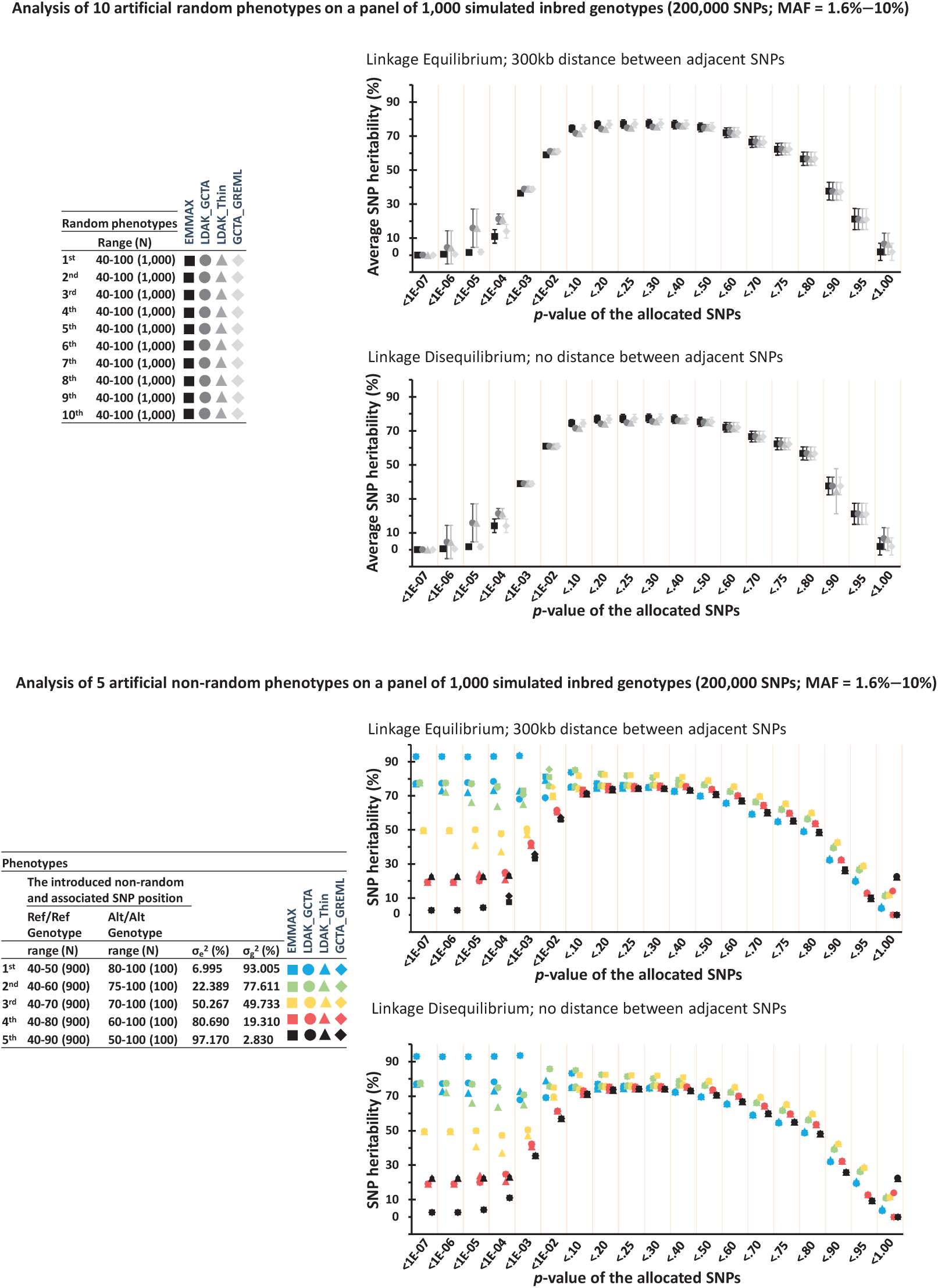
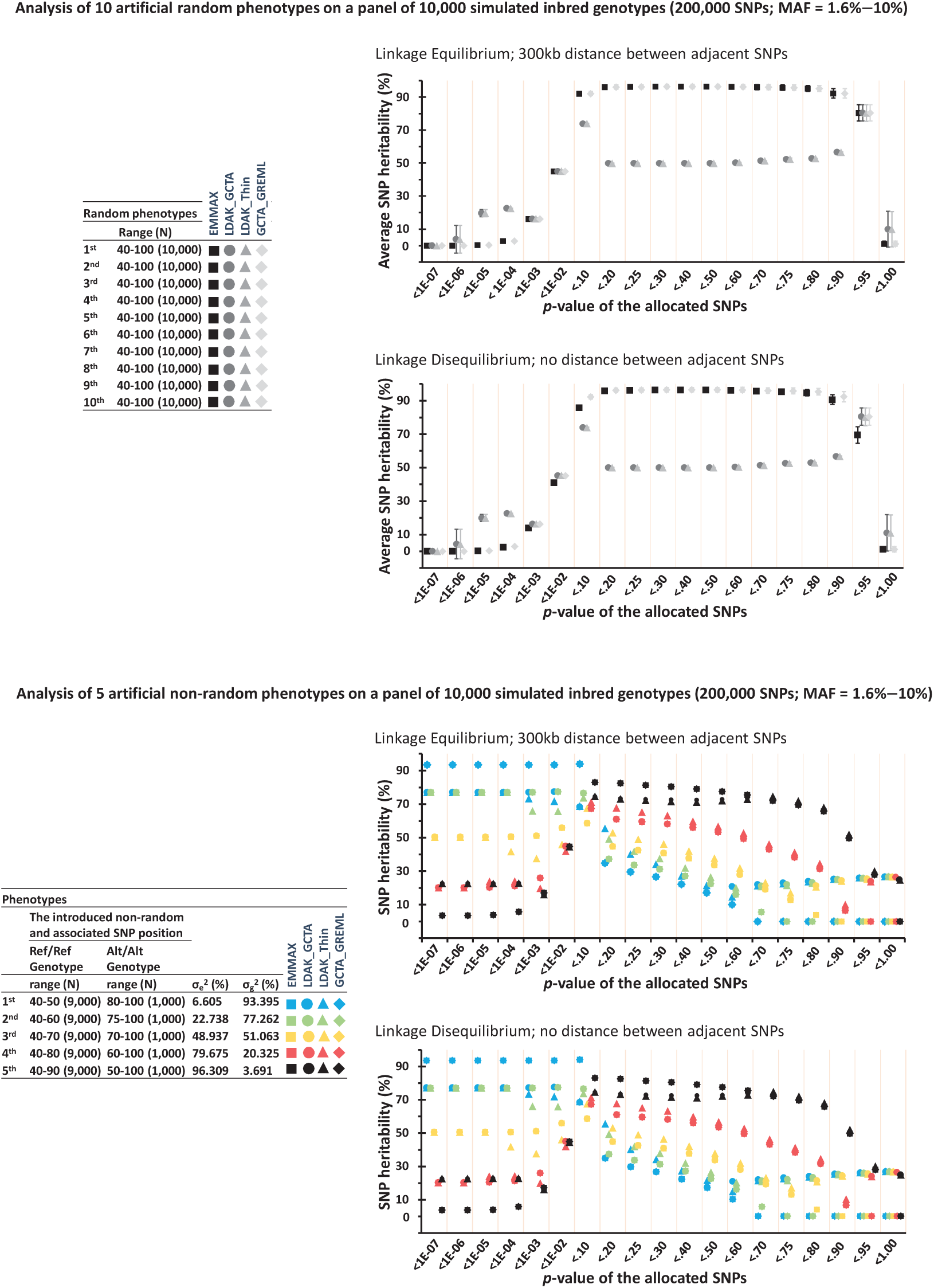
The SNP heritability profiles for simulated quantitative-phenotypes and SNPs were analyzed comparing linkage disequilibrium to linkage equilibrium. The analysis compares linkage disequilibrium (i.e., 200k SNPs on a 200kb chromosome) and linkage equilibrium (i.e., 300kb distance between adjacent SNPs). Initial GWAS was performed using EMMAX to realize association levels for the SNPs. The four different approaches of EMMAX, LDAKGCTA, LDAK-Thin, and GCTA-GREML were then applied to estimate SNP heritability for the clusters of SNP hits captured using the association p-values. N represents the sample sizes, i.e., number of genotypes. MAF stands for the minor allele frequency. The average SNP heritability estimates were obtained for 10 independent random phenotypes within identical ranges. The error bars represent 99% confidence intervals. The SNP heritability estimates were obtained for five artificial non-random phenotypes which were associated with a simulated non-random SNP position introduced within the simulated genotypes. By ensuring additive-only effects, the non-random phenotypes were defined at different quantitative ranges among the genotypic groups of the introduced associated SNP position, i.e., Ref/Ref and Alt/Alt for the simulated inbred genotypes. The intragenotype assignment of the phenotypes, while being within their specific ranges, was still random. The inter-genotype variances of the phenotypes are shown as σ_g_^2^. σ_e_ ^2^ represents the intra-genotype variations. The analyses of inbred genotypes using GCTA-GREML were performed based on the “inbred” algorithm of GCTA.

**Supplementary Fig. 6:**
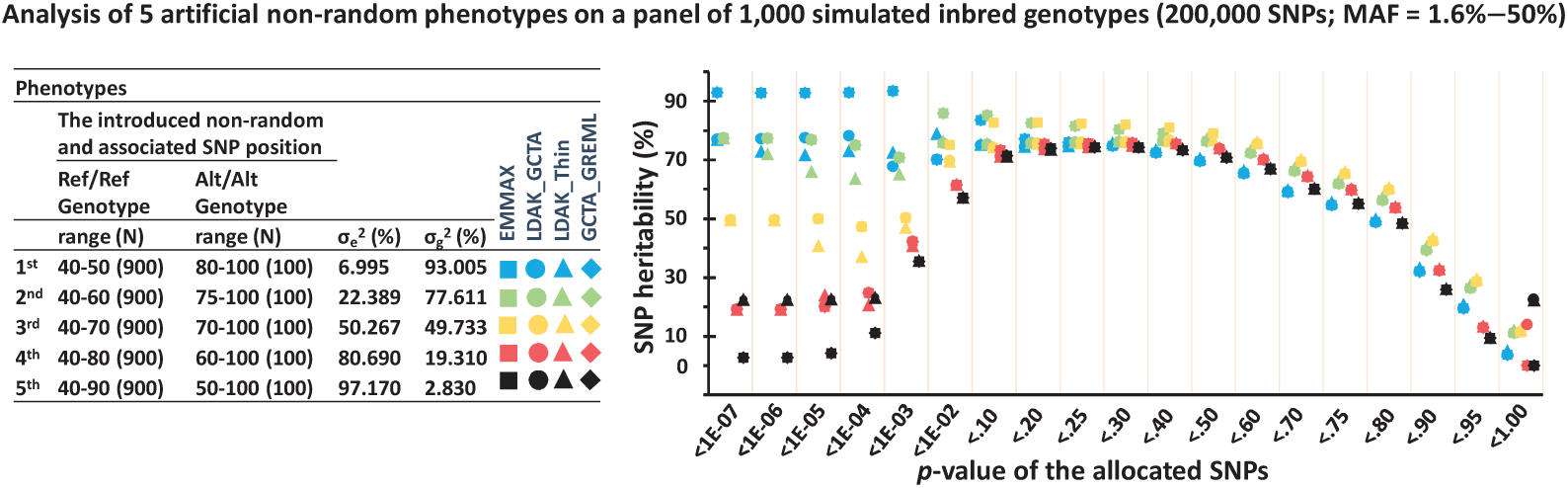
The SNP heritability profiles for simulated quantitative-phenotypes and SNPs with minor allele frequencies ranging from 1.6%–50%. Initial GWAS was performed using EMMAX to realize association levels for the SNPs. The four different approaches of EMMAX, LDAKGCTA, LDAK-Thin, and GCTA-GREML were then applied to estimate SNP heritability for the clusters of SNP hits captured using the association *p-*values. N represents the sample sizes, i.e., number of genotypes. MAF stands for the minor allele frequency. The SNP heritability estimates were obtained for five artificial non-random phenotypes which were associated with a simulated non-random SNP position introduced within the simulated genotypes. By ensuring additive-only effects, the non-random phenotypes were defined at different quantitative ranges among the genotypic groups of the introduced associated SNP position, i.e., Ref/Ref and Alt/Alt for the simulated inbred genotypes. The intragenotype assignment of the phenotypes, while being within their specific ranges, was still random. The inter-genotype variances of the phenotypes are shown as σ_g_^2^. σ_e_^2^ represents the intra-genotype variations. The analyses of inbred genotypes using GCTA-GREML were performed based on the “inbred” algorithm of GCTA.

**Supplementary Fig. 7:**
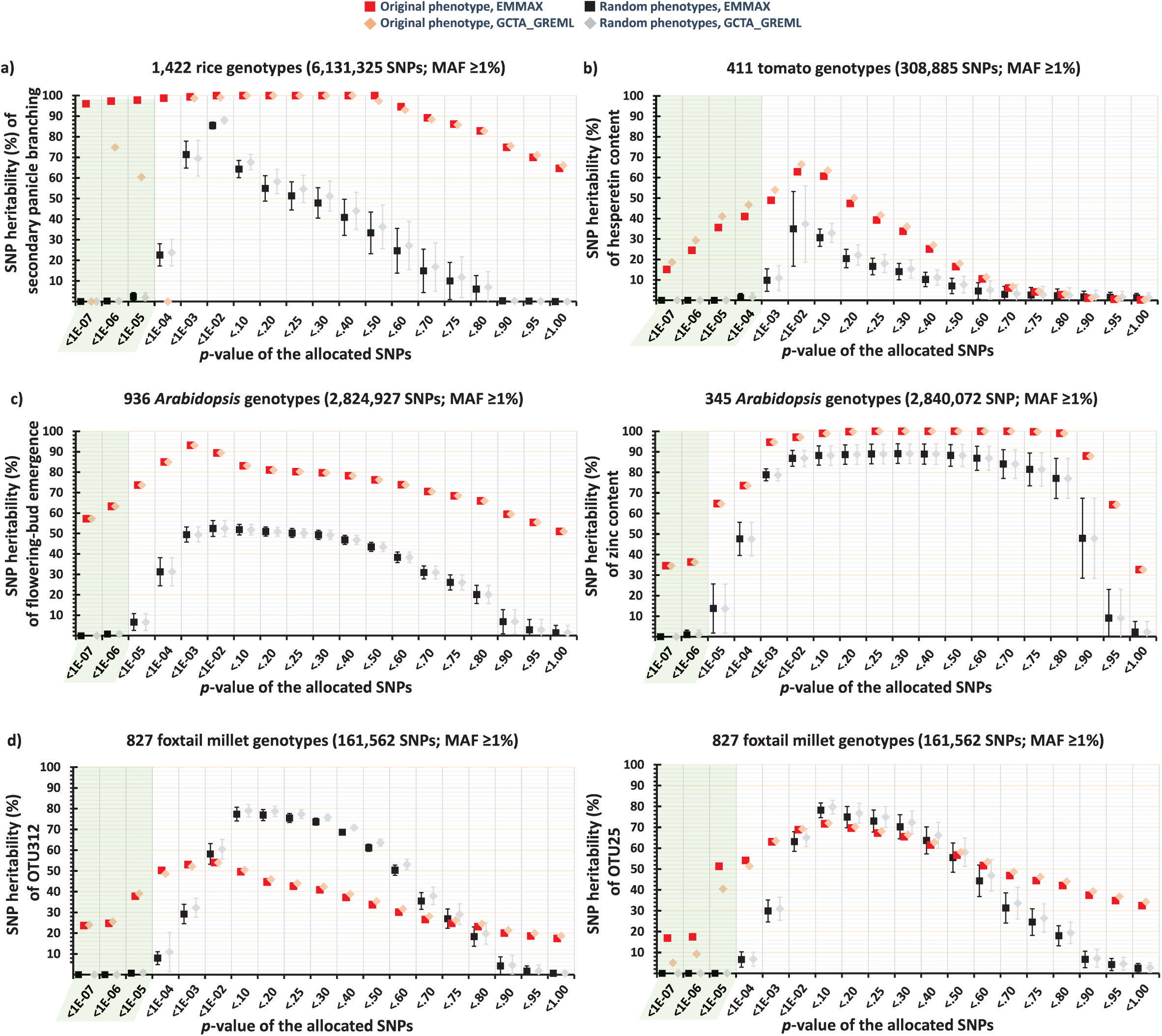
SAFE-h^2^ estimates SNP heritability by minimizing the contributions from false-positive SNP hits, a-d) The safe-intervals for p-value are highlighted in light green representing the area with minimum contributions to heritability by false-positive SNP hits. The two different approaches of EMMAX and GCTA-GREML were applied to estimate the SNP heritability for the clusters of SNP hits captured using the association p-values. MAF stands for the minor allele frequency. The average estimations are illustrated for six independent random phenotypes (simulated within the range of original phenotype) and the error bars represent standard deviations among the six estimations. GCTA-GREML was performed based on the "inbred" algorithm. OUT: Operational Taxonomic Unit.

**Supplementary Fig. 8:**
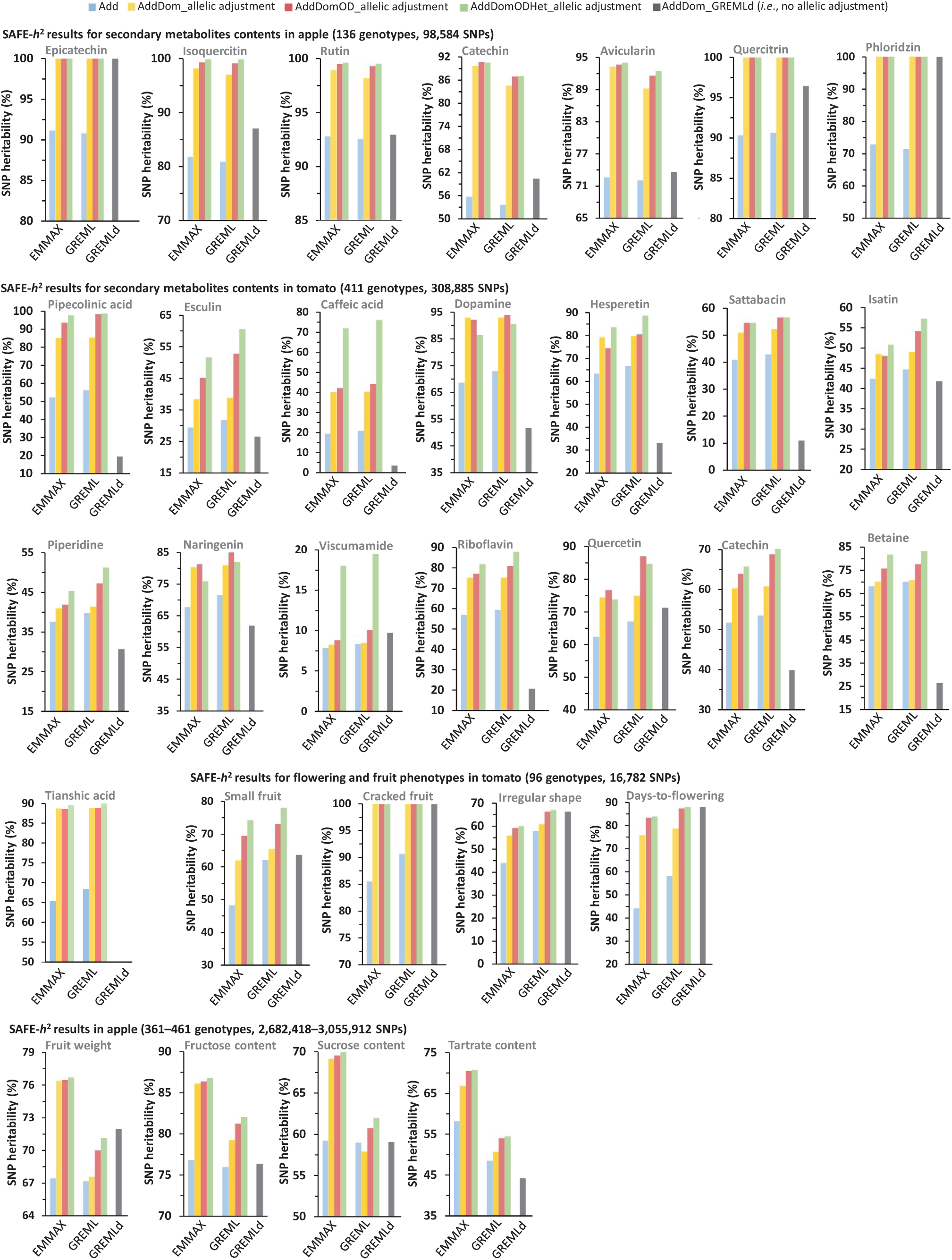
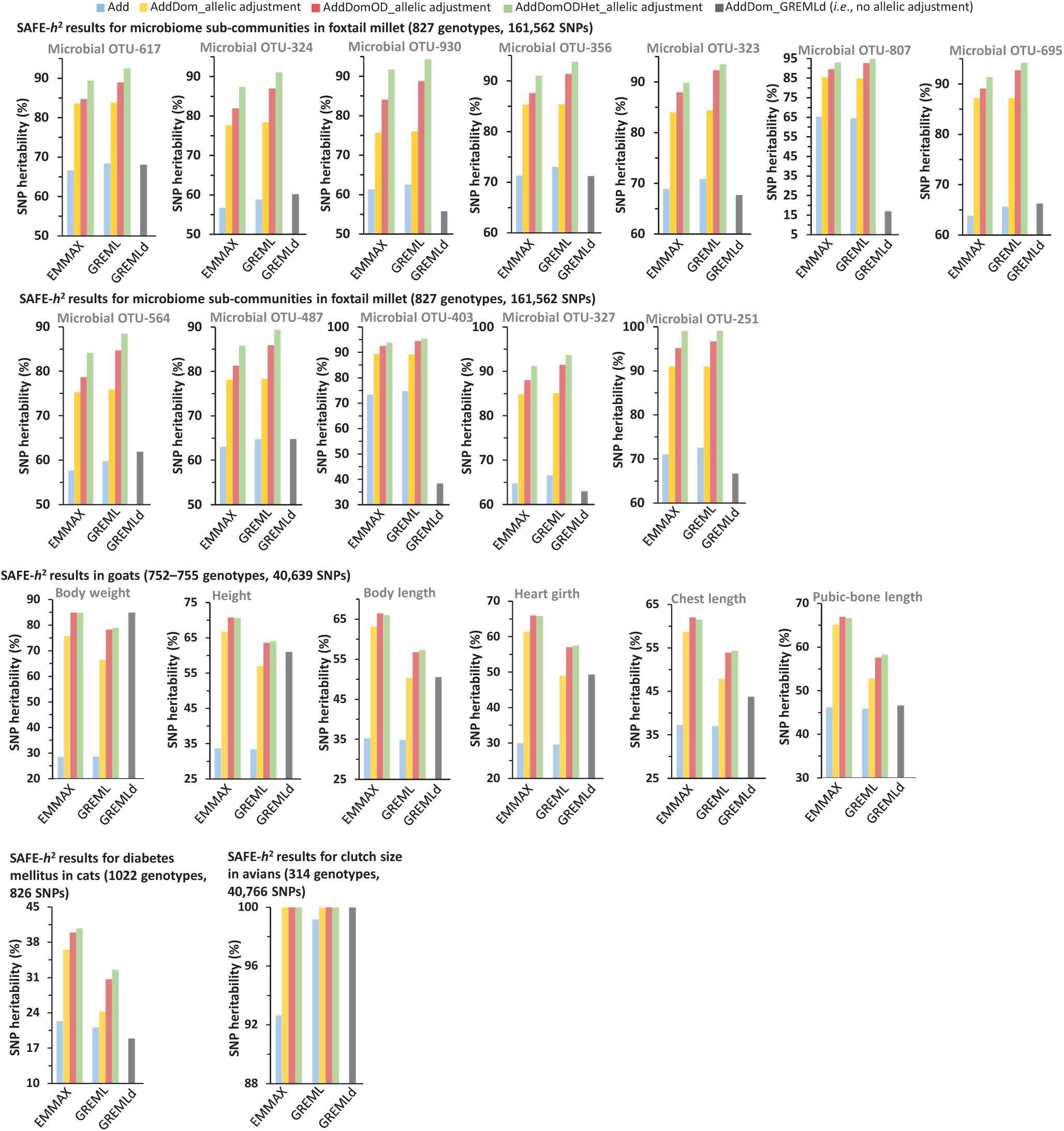
The SAFE-*h*^2^ estimates of SNP heritability using additive-only model and the combined additive and non-additive models, compared to GREMLd estimates. The analyses of tomato and foxtail millet genotypes using GCTA-GREML/GREMLd models were performed based on the “inbred” algorithm of GCTA. The average inbreeding coefficients (Fhat) of tomato and foxtail millet genotypes were > 0.8. Add: heritability based on additive effects, AddDom: heritability based on additive and dominance effects, AddDomOD: heritability based on additive, dominance and overdominance effects, AddDomODHet: heritability based on additive, dominance, overdominance, and heterosis-like effects, GREMLd: heritability estimated by GREMLd model on additive and dominance effects (*i.e*., without allelic adjustment), MAF: minor allele frequency, OUT: Operational Taxonomic Unit. MAF was ≥ 1% in all the analyses.

**Supplementary Fig. 9:**
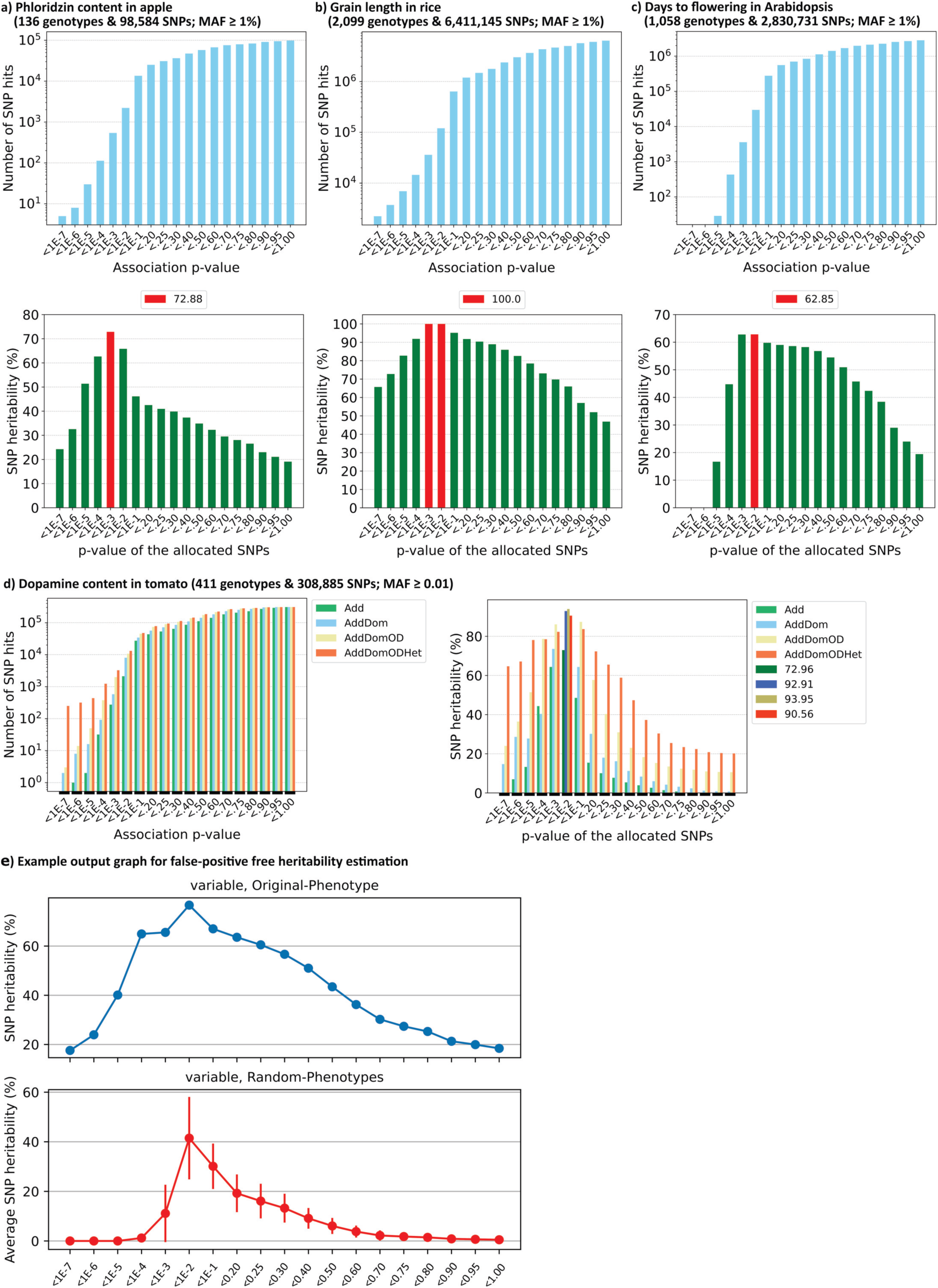
SAFE-*h*^2^ generates bar graphs for both the *p*-value-guided SNP clusters and the SNP heritability profiles. The SNP heritability estimates are highlighted by red color in (a), (b), (c), and by darker colors in (d). e) Bars represent the standard deviations among six random phenotypes. Add: heritability based on additive effects. AddDom: heritability based on additive and dominance effects. AddDomOD: heritability based on additive, dominance and overdominance effects. AddDomODHet: heritability based on additive, dominance, overdominance, and heterosis-like effects. MAF: minor allele frequency.

**Supplementary Table 1.**
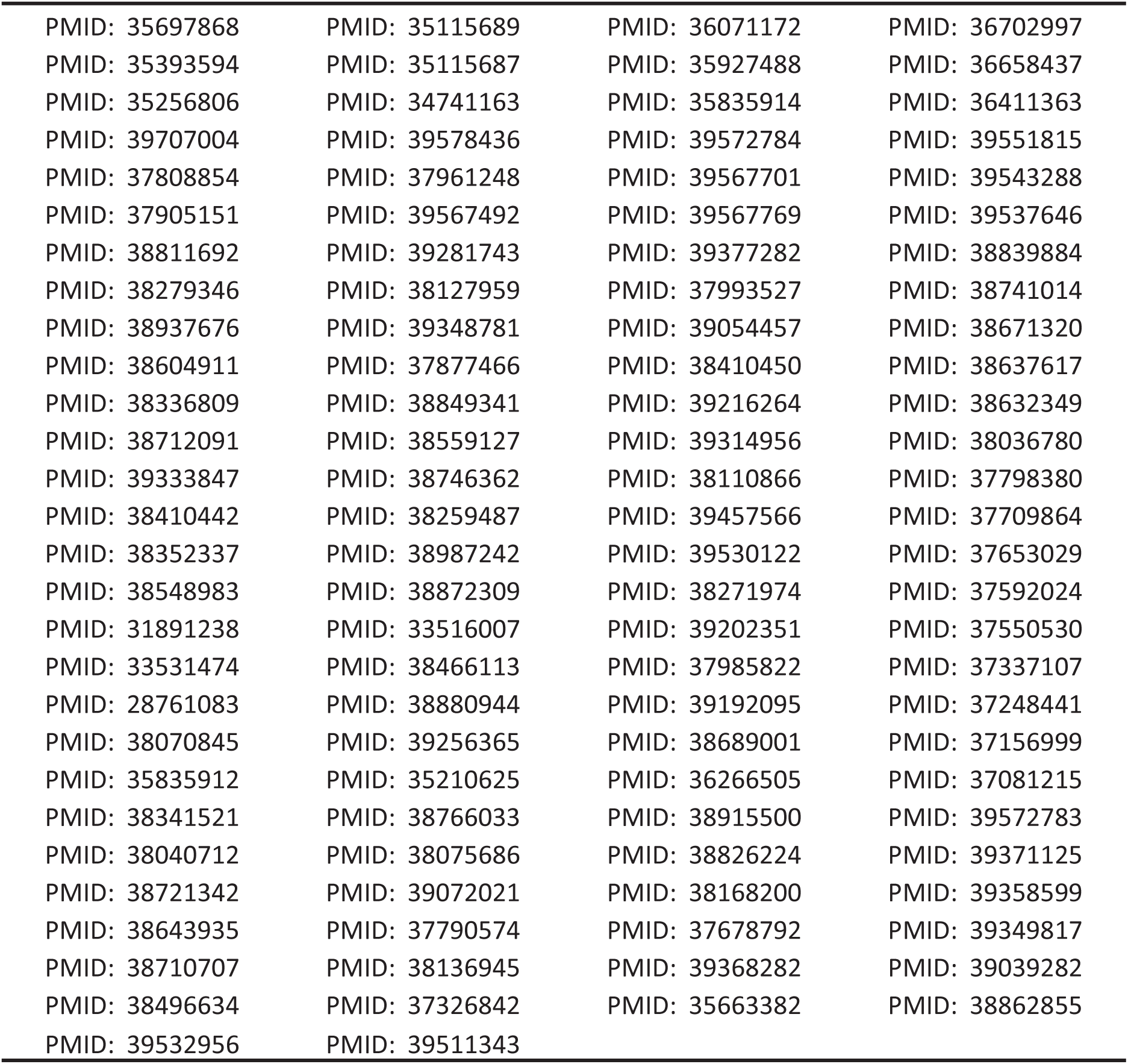
The PubMed PMID accessions of the recently published studies used to extract SNP heritability estimates for different traits and species.

**Supplementary Table 2.**
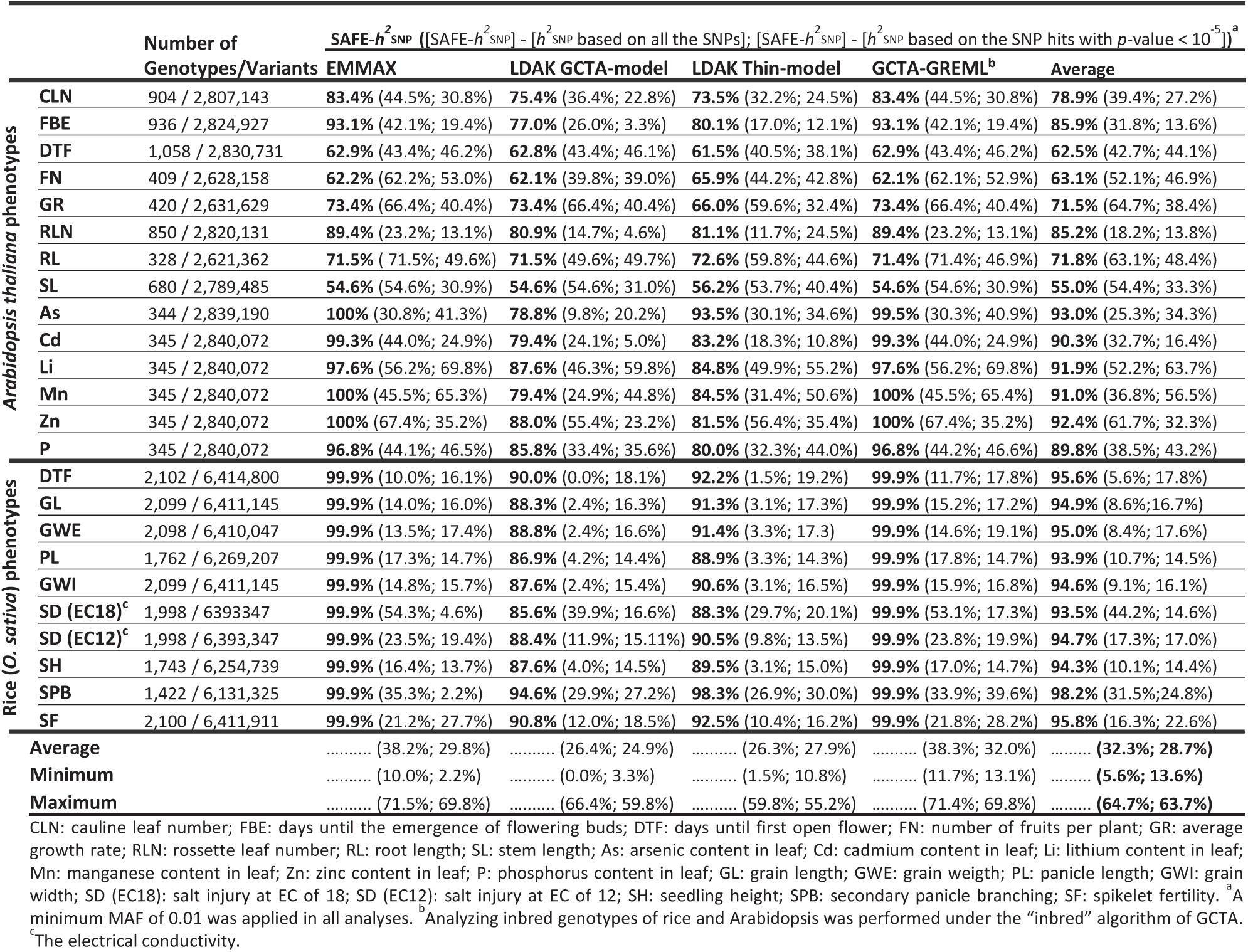
SAFE-*h*^2^ improves the heritability estimations for 24 traits in *Arabidopsis* and rice by preventing the negative contributions of weakly-associated genomic regions.

**Supplementary Table 3.**
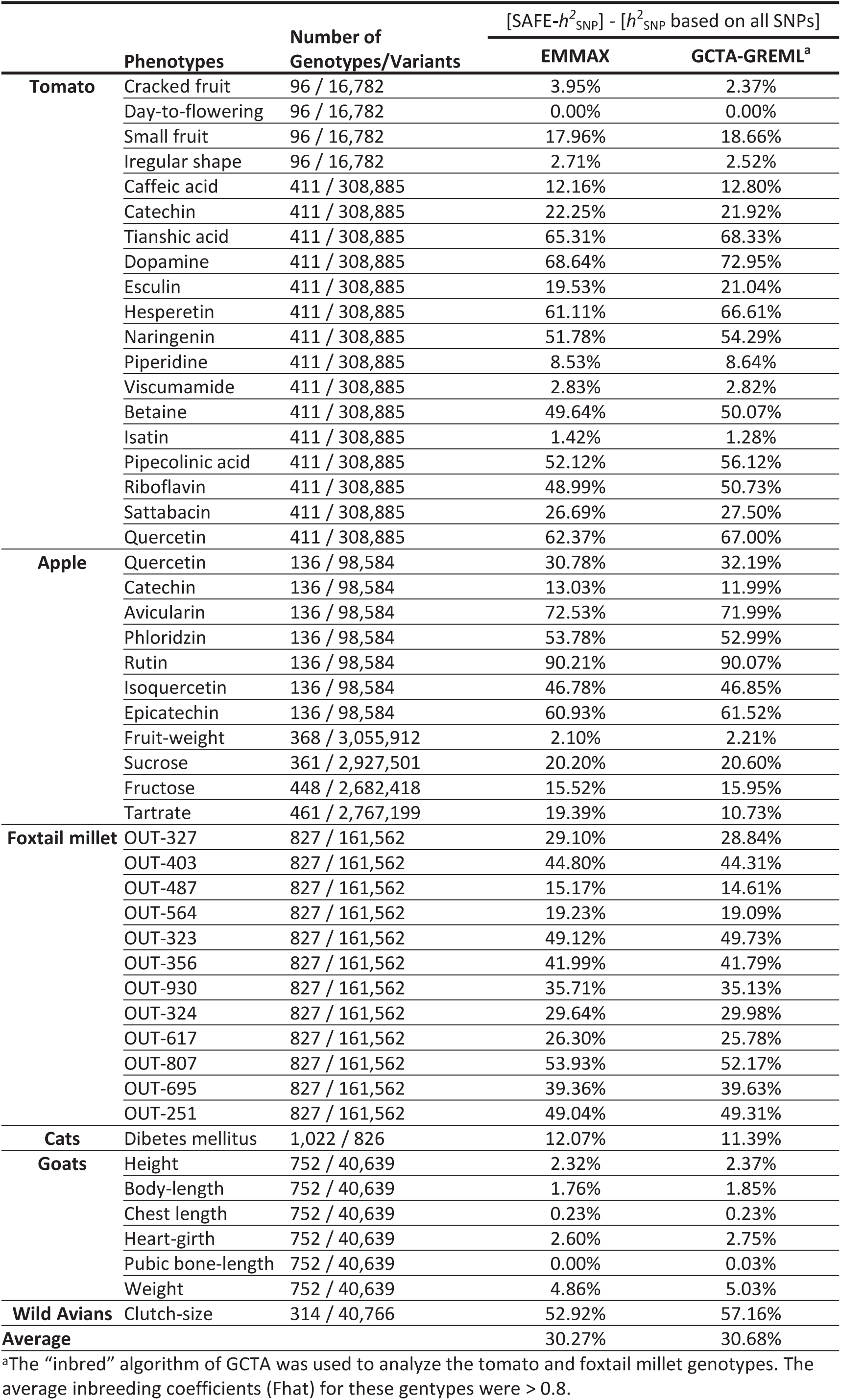
Heritability profiling by SAFE-*h*^2^ reveals negative contributions of SNPs with large association *p*-values by examining 50 phenotypes in six species.

**Supplementary Table 4:** BayesR SNP heritability estimates for simulated quantitative-phenotypes using the 200K simulated SNPs in LE and LD.

| The introduced non-random (associated) SNP position |  |  |  |  | On a 200kb chromosome (Linkage Disequilibrium) |  | 300kb distance between two adjacent SNPs (Linkage Equilibrium) |  |  |  |
| --- | --- | --- | --- | --- | --- | --- | --- | --- | --- | --- |
| Ref/Ref Genotype | | Alt/Alt Genotype | | | BayesR | | Process time (BayesR/SAFE-$h^2$) | BayesR | | Process time (BayesR/SAFE-$h^2$) |
| Phenotypes | range (N) | range (N) | $\sigma_e^2$ (%) | $\sigma_g^2$ (%) | $h^2$ (%) | Process time | | $h^2$ (%) | Process time | |
| 1<sup>st</sup> | 40-50 (900) | 80-100 (100) | 6.995 | 93.005 | na | 150 min | 10.7 | na | 150 min | 10.7 |
| 2<sup>nd</sup> | 40-60 (900) | 75-100 (100) | 22.389 | 77.611 | na | 150 min | 10.7 | na | 150 min | 10.7 |
| 3<sup>rd</sup> | 40-70 (900) | 70-100 (100) | 50.267 | 49.733 | na | 150 min | 10.7 | 34.67 | 150 min | 10.7 |
| 4<sup>th</sup> | 40-80 (900) | 60-100 (100) | 80.69 | 19.31 | 6.04 | 150 min | 10.7 | 8.33 | 150 min | 10.7 |
| 5<sup>th</sup> | 40-90 (900) | 50-100 (100) | 97.17 | 2.83 | 3.47 | 150 min | 10.7 | 1.07 | 150 min | 10.7 |

Ref/Ref and Alt/Alt for the introduced associated single SNP, N: Number of genotypes (individuals)
$\sigma_g^2$ : Inter-genotype variances of the phenotypes
$\sigma_e^2$ : Intra-genotype variations

**Supplementary Table 5:** BayesR SNP heritability estimates for simulated quantitative-phenotypes using the chr. 22 SNPs from human HapMap phase 1 genotypes.

| The introduced non-random (associated) SNP position |  |  |  |  |  | Real genotypes based on Chr22 + introduced single associated SNP |  |  |
| --- | --- | --- | --- | --- | --- | --- | --- | --- |
| Ref/Ref Genotype | | Ref/Alt Genotype | Alt/Alt Genotype | | | BayesR | | Process time (BayesR/SAFE-$h^2$) |
| Phenotypes | range (N) | range (N) | range (N) | $\sigma_e^2$ (%) | $\sigma_g^2$ (%) | $h^2$ (%) | Process time | |
| 1<sup>st</sup> | 40-50 (982) | 80-100 (103) | 120-150 (7) | 5.704 | 94.296 | na | 120 min | 8.6 |
| 2<sup>nd</sup> | 40-60 (982) | 75-100 (103) | 112-150 (7) | 21.037 | 78.963 | na | 120 min | 8.6 |
| 3<sup>rd</sup> | 40-70 (982) | 70-100 (103) | 105-150 (7) | 41.501 | 58.499 | na | 120 min | 8.6 |
| 4<sup>th</sup> | 40-80 (982) | 60-100 (103) | 90-150 (7) | 72.099 | 27.901 | 22.21 | 120 min | 8.6 |
| 5<sup>th</sup> | 40-90 (982) | 50-100 (103) | 75-150 (7) | 91.416 | 8.584 | 6.17 | 120 min | 8.6 |

Ref/Ref, Ref/Alt, and Alt/Alt for the introduced associated single SNP
$\sigma_g^2$ : Inter-genotype variances of the phenotypes
$\sigma_e^2$ : Intra-genotype variations
N: number of genotypes (individuals)

**Supplementary Table 6:**
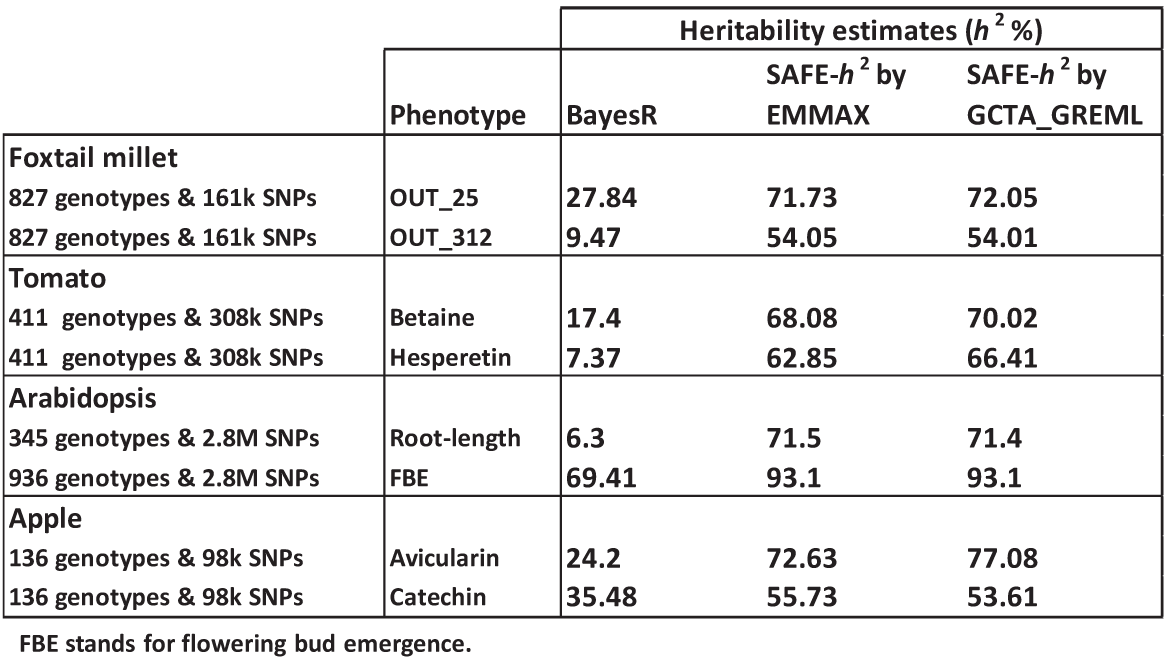
The SNP heritability estimates for real phenotypes using BayesR and SAFE-*h*^2^.

**Supplementary Table 7.**
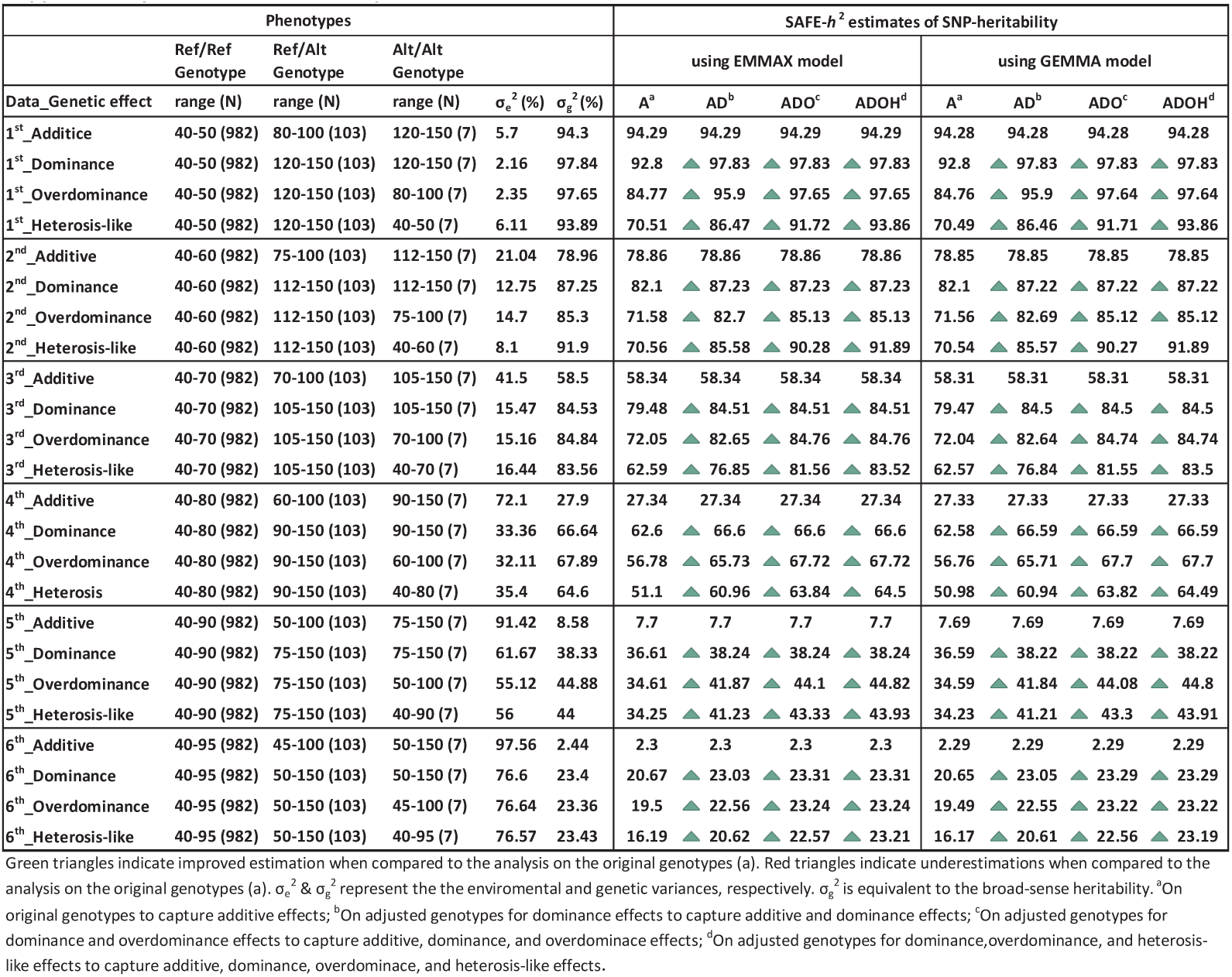
SNP heritability estimations on additive or additive + non-additive effects of a biallelic locus.

**Supplementary Table 8.** Heritability estimates based on p-values, calculated using EMMAX and GEMMA.

| | | Heritability estimates ($h^2$ %) | | | | | | | | | | | |
| --- | --- | --- | --- | --- | --- | --- | --- | --- | --- | --- | --- | --- | --- |
| | | Association analysis using EMMAX ($p$-value source) | | | | | | Association analysis using GEMMA ($p$-value source) | | | | | |
| | Phenotype | EMMAX (on all SNPs) | SAFE-$h^2$ using EMMAX | GCTA_GREML (on all SNPs) | SAFE-$h^2$ using GCTA_GREML | GEMMA (on all SNPs) | SAFE-$h^2$ using GEMMA | EMMAX (on all SNPs) | SAFE-$h^2$ using EMMAX | GCTA_GREML (on all SNPs) | SAFE-$h^2$ using GCTA_GREML | GEMMA (on all SNPs) | SAFE-$h^2$ using GEMMA |
| Foxtail millet |  |  |  |  |  |  |  |  |  |  |  |  |  |
| 827 genotypes & 161k SNPs | OUT_25 | 32.42 | 71.73 | 34.26 | 72.05 | 32.94 | 74.41 | 32.43 | 94.97 | 34.26 | 95.17 | 32.94 | 97.97 |
| 827 genotypes & 161k SNPs | OUT_312 | 17.48 | 54.05 | 18.68 | 54.01 | 17.86 | 57.37 | 17.49 | 85.01 | 18.68 | 86.02 | 17.86 | 88.567 |
| Tomato |  |  |  |  |  |  |  |  |  |  |  |  |  |
| 411 genotypes & 308k SNPs | Betaine | 18.43 | 68.08 | 19.99 | 70.02 | 18.02 | 67.35 | 18.43 | 73.23 | 19.99 | 74.85 | 18.02 | 74.46 |
| 411 genotypes & 308k SNPs | Hesperetin | 0.21 | 63.39 | 0.23 | 66.84 | 0.26 | 62.5 | 0.21 | 62.04 | 0.23 | 65.58 | 0.26 | 61.18 |
| Arabidopsis |  |  |  |  |  |  |  |  |  |  |  |  |  |
| 345 genotypes & 2.8M SNPs | Root-length | 0.01 | 71.47 | 0.01 | 71.42 | 0 | 71.41 | 0.01 | 71.58 | 0 | 71.52 | 0 | 71.52 |
| 936 genotypes & 2.8M SNPs | FBE | 51.03 | 93.14 | 51 | 93.13 | 51 | 93.13 | 51.03 | 93.58 | 51 | 93.58 | 51 | 93.58 |
| Apple |  |  |  |  |  |  |  |  |  |  |  |  |  |
| 136 genotypes & 98k SNPs | Avicularin | 0.09 | 72.63 | 0.09 | 72.08 | 0.01 | 80.34 | 0.09 | 79.74 | 0.09 | 79.26 | 1.47 | 86.54 |
| 136 genotypes & 98k SNPs | Catechin | 42.69 | 55.73 | 41.61 | 53.61 | 45.05 | 58.88 | 42.69 | 95.77 | 41.61 | 95.65 | 45.05 | 97.09 |

### Supplementary text 1: Independently simulated datasets cannot be used as replicates

When simulating datasets, every dataset shows the similar downward bias in heritability estimates by adding unassociated SNPs, i.e., with large association *p*-values (see Fig. 2b and Suppl. Fig. 2-6). This downward bias is also observed in real-datasets (see Fig 2c, Suppl. Fig. 7, Suppl. Fig. 9, Suppl. Table 2 and 3).

We can use *p*-values from one simulated dataset to make heritability estimates in another simulated dataset (as a replicated study), but only if the second dataset is not independently simulated from scratch. Under these conditions, we will observe the same pattern of downward bias shown in Fig. 2b and Suppl. Figs. 2-6.

An example of a proper replication:

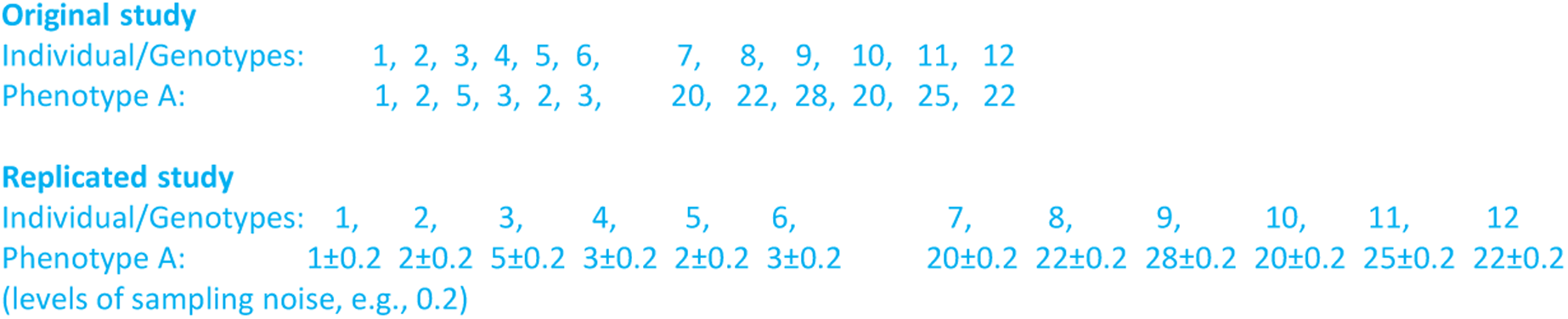

In contrast, it is not possible to have replication through full re-simulation of either of genotypes or phenotypes. When re-simulating phenotype from scratch, we generate new independent phenotypes, for example the original was phenotype A and now we have observation of phenotype B.

An example of a wrong replication:

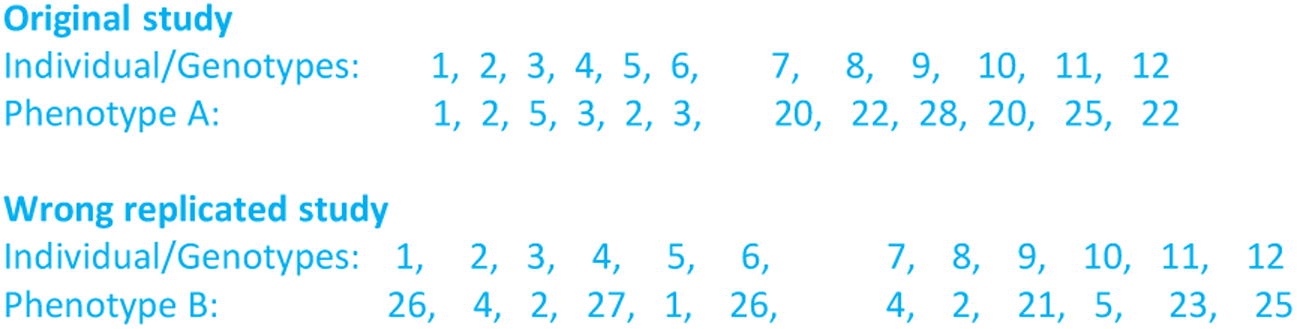

Here, we cannot use the *p*-values from the original dataset to make heritability estimation in the second dataset which was simulated independently from scratch (i.e., wrong replicated study).

When re-simulating genotypes, it also means a complete new cohort, for example the original was a cohort on disease A and the re-simulated is a cohort on disease B. So, that is why *p*-values obtain from one simulation, so-called “original study/cohort”, cannot be used in another dataset which is simulated from scratch.

## Notes

https://github.com/SAFE-h2/SAFE-h2

https://figshare.com/s/7193d332894f11deb1a5

### Competing Interest Statement

The authors have declared no competing interest.

### Summary of Updates

A description is provided on how simulated datasets can be used for replicated analysis, along with supplementary text. Additionally, minor edits have been made to the text.

